# Attenuated gaze synchrony in autism and its non-additive modification by co-occurring ADHD: a transdiagnostic eye-tracking study of naturalistic viewing

**DOI:** 10.64898/2026.05.11.724367

**Authors:** Xin Di, Bharat B. Biswal

**Author notes:** Corresponding author: 604 Fenster Hall, University Heights Newark, NJ, 07102, USA.

## Abstract

**Background:** Autism spectrum disorder and attention-deficit/hyperactivity disorder co-occur frequently, but whether co-occurrence compounds or alters the visual-attentional differences associated with autism remains unresolved. Inter-subject correlation (ISC) during naturalistic viewing provides a model-free index of how closely an individual’s gaze and pupillary responses track those of typically developing peers.

**Methods:** We analyzed eye-tracking data from the Healthy Brain Network across four videos differing in social and visual content (N = 2476 children and adolescents; 1186–2042 per video). Gaze and pupil ISC were computed against a typically developing reference, gaze as a two-dimensional complex correlation. Dimensional models used parent-reported autistic (SRS-2) and ADHD (SWAN) traits; categorical models used a 2 × 2 (ASD × ADHD) analysis of covariance. All models adjusted for age, sex, tracker configuration and screen shape.

**Results:** Autistic traits predicted reduced gaze synchrony in three of four videos, independently of ADHD traits. An autism diagnosis was associated with reduced gaze synchrony in all four videos (β = −0.78 to −0.45 SD; q ≤ 0.009). The ASD × ADHD interaction was significant and positive in three videos (β = 0.46–0.65; lower 95% confidence limits ≥ 0.16; q ≤ 0.005), indicating that the reduction associated with autism was smaller when ADHD co-occurred. The interaction was significant in all four videos within a single acquisition configuration and was robust to adjustment for co-occurring psychopathology and same-day stimulant status and to down-weighting of extreme observations. Ten stimulus-independent oculomotor measures from the same recordings showed no consistent group differences. Absolute gaze phase did not survive comparison with a magnitude-matched null.

**Limitations:** The group with autism and without ADHD was small (45–78 children per video), so the interaction is estimated imprecisely. The findings were not tested in an independent cohort. Diagnoses were made without standard autism research instruments, eye-tracker settings varied, medication status was mostly unknown, and the design was cross-sectional.

**Conclusions:** Co-occurring ADHD is associated with a smaller, not larger, reduction in gaze synchrony than autism alone, which argues against additive models of their co-occurrence. Inter-subject synchrony captures group structure that coarse summaries of eye movement do not.

## 1. Background

When a group of children watches the same film, their eyes move together. A character turns and gaze follows; a face enters the frame and gaze settles on it; an object falls and gaze drops to meet it. No one instructs them to look at these things. The film itself does the work, and the result is that at most moments most children are looking at nearly the same point on the screen.

Children differ in how closely they follow. Some track the crowd almost exactly. Others spend the same minutes looking at the edges of the frame, at background objects, or at whatever happens to have moved last. These differences are stable enough to be measured, and they are pronounced in autism. Autistic children look less at faces and eyes and more at bodies and objects when viewing social scenes (Klin et al., 2002), fixate more idiosyncratically than their peers (Falck-Ytter et al., 2013), and do so from the first years of life, reliably enough that the pattern now underpins a diagnostic device cleared for clinical use (Jones et al., 2023). Attention-deficit/hyperactivity disorder also alters how the eyes move, though differently: children with ADHD shift gaze more frequently and hold fixation less steadily (Munoz et al., 2003). During naturalistic viewing, their gaze also corresponds less closely to the locations a model of typical viewing predicts (Dziemian et al., 2025).

These two conditions co-occur far more often than chance allows: ADHD is among the most common co-occurring diagnoses in autism, affecting roughly a quarter to a third of autistic people (Lai et al., 2019), and the two share genetic and phenotypic variance (Ronald et al., 2008). What remains unsettled is what the combination does. The intuitive expectation is that difficulties add: a child with both conditions should look less like the crowd than a child with either alone. The alternative is that the two alter visual behavior in ways that do not simply sum. Frequent gaze shifting and prolonged idiosyncratic fixation, occurring in the same child, may produce something other than more of each.

Recent large-scale work bears on this question without resolving it. Resting-state connectivity shows separable, additive signatures for autistic and attentional traits with no interaction between them (Norman et al., 2025), and discrete behavioral assays in the same cohort likewise show none (Silvan et al., 2026). Both paradigms, however, measure something other than sustained engagement with a continuous stimulus. A child at rest is not watching anything, and a child performing a structured task is doing what the task demands.

Naturalistic viewing offers a measure the others do not. When participants watch the same film, the degree to which an individual’s moment-to-moment response matches the group response can be computed directly, without specifying in advance which features of the film should drive attention. This quantity, inter-subject correlation, was developed for brain imaging (Hasson et al., 2004; Nastase et al., 2019) and has since been applied to gaze itself, where reduced similarity to normative viewing patterns has been reported in autism (Avni et al., 2020). Gaze is a particularly direct substrate for it: the measure asks whether a child is looking where other children look, which is a description of attention rather than an inference about it.

Applying inter-subject correlation to gaze raises a question that does not arise for a single time series. Gaze is two-dimensional, and treating the horizontal and vertical axes as independent correlations discards the relationship between them. Mapping the two axes into a single complex-valued trajectory preserves that relationship and yields a correlation with two components: a magnitude, indexing how strongly a child’s gaze is entrained to the group, and a phase, indexing any rotation of the child’s gaze trajectory relative to the group’s. Whether the second component carries information the first does not is an empirical question, and one we address directly.

Pupil size provides a second, independent channel. Pupillary fluctuations during viewing track arousal and attentional engagement (Aston-Jones & Cohen, 2005), and the same synchrony logic applies: a child whose pupil dilates when other children’s pupils dilate is responding to the film as they are.

We analyzed eye-tracking data from 2476 children and adolescents in the

Healthy Brain Network (Alexander et al., 2017), a transdiagnostic cohort in which autism, ADHD and their co-occurrence are all well represented, recorded while they watched four short videos. The videos differ in what they ask of a viewer: one follows a boy and a three-legged puppy through a wordless emotional exchange, another a geometric pattern unfolding to narration. If reduced synchrony in autism reflects diminished engagement with social content specifically, it should be confined to the former. We asked whether autistic and ADHD traits predict reduced gaze and pupillary synchrony, whether their clinical co-occurrence produces an additive or a non-additive effect, and whether any effect we observed could be recovered from coarse oculomotor measures of the same recordings.

## 2. Methods

### 2.1. Participants and Clinical Assessment

Data were obtained from the Healthy Brain Network (HBN; (Alexander et al., 2017)), Releases 1–11. Participants with eye-tracking data passing quality control for at least one of four naturalistic viewing paradigms and with available clinical characterization were included, yielding a final sample of 2476 (1565 male, 911 female; mean age 10.39 years, SD 3.44). Guardians or participants of legal age provided written informed consent; study protocols were approved by the Chesapeake Institutional Review Board.

Consensus DSM-5 clinical diagnoses were established by licensed clinicians using the Kiddie Schedule for Affective Disorders and Schizophrenia, Computerized version (K-SADS-COMP; (Townsend et al., 2020)), a semi-structured interview incorporating child and caregiver reports. Gold-standard autism-specific instruments (ADOS-2, ADI-R) were not available for this sample; this constrains interpretation of the categorical analyses and is addressed in the Limitations. For categorical models, participants were stratified into a strict 2 × 2 factorial design: typically developing (TD) controls (n = 239), ADHD without ASD (n = 1156), ASD without ADHD (n = 99), and comorbid ASD+ADHD (n = 267). A heterogeneous “Other” psychiatric group (n = 715; e.g., anxiety or learning disorders without ASD or ADHD) was excluded from the categorical analyses and retained in the dimensional analyses (Section 2.7.1).

Continuous neurodevelopmental traits were quantified using parent reports on the Social Responsiveness Scale, Second Edition (SRS-2, hereafter SRS; (Constantino, 2021)) and the Strengths and Weaknesses of ADHD Symptoms and Normal Behavior scale (SWAN; (Swanson et al., 2012)), on both of which higher scores indicate greater symptom severity. To harmonize data across age-specific versions, SRS-2 Total T-scores were used for both Preschool and School-Age forms. SRS subscale scores, full-scale IQ (Wechsler Intelligence Scale for Children), Child Behavior Checklist (CBCL, (Achenbach & Rescorla, 2023)) Total T-scores, and clinician-recorded anxiety and oppositional/conduct diagnoses were retained for severity and ascertainment analyses. Acute medication status on the day of assessment was documented using the HBN DailyMeds instrument for a subset of participants (n = 514, 20.8%); medication status is unknown for the remainder and is not treated as unmedicated.

No prior power calculation was performed; the sample comprised all participants with usable recordings.

### 2.2. Naturalistic Viewing Stimuli

Participants underwent a task-free naturalistic viewing paradigm in a sound-shielded room as part of a combined electroencephalography and eye-tracking session, seated facing a computer display; screen geometry and viewing distance varied across recordings and are addressed in Sections 2.3 and 2.4. The paradigm featured four videos that differ in social complexity, emotional valence, and visual salience:

- ***The Present* (TP; 3:22):** An emotionally complex, character-driven animated short heavily reliant on subtle social cues.
- ***Despicable Me* (DM; 2:50):** A fast-paced, visually dynamic clip from an animated comedy.
- ***Diary of a Wimpy Kid* (WK; 1:57):** A live-action movie trailer featuring school-based social dynamics.
- ***Fun with Fractals* (FF; 2:43):** An abstract, non-social educational video featuring repetitive geometric patterns.

No explicit cognitive or behavioral task was required during viewing. The HBN protocol documentation states that *Despicable Me*, *Diary of a Wimpy Kid* and *Fun with Fractals* were presented in randomized order and that *The Present* was always presented last; we did not verify presentation order independently, and given the discrepancies between that documentation and the recordings described in Section 2.3, it should be regarded as nominal.

### 2.3. Eye-Tracking and Pupillometry Acquisition

Electroencephalography was recorded simultaneously but is not analyzed here. Published HBN protocol documentation describes a single infrared eye tracker (iView-X RED-m, SensoMotoric Instruments GmbH) sampling at 120 Hz with 0.1° spatial resolution and 0.5° gaze accuracy (Alexander et al., 2017). Rather than assume a uniform acquisition setup, we parsed the metadata header of every raw file and tabulated the configuration each recording was actually produced under. Among the 8233 recordings that passed gaze preprocessing (Section 2.4), three nominal sampling rates were present: 30 Hz (n = 1038 recordings), 60 Hz (n = 5525), and 120 Hz (n = 1670). A corneal-reflection (CR) correction stage, which yields a calibrated pupil diameter in millimeters, was active in every 30 Hz file and in none of the 120 Hz files; at 60 Hz it was present in 4699 recordings and absent in 826. The calibrated display area was likewise not fixed: 800 × 600 px in 5416 recordings, 1440 × 900 px in 2742, and 1600 × 900 px in 75. Physical stimulus dimensions and head distance were also recorded per file.

We were unable to reconcile this heterogeneity with the single-system protocol description from publicly available documentation. A plausible explanation is that hardware, firmware, or export settings changed over the course of an ongoing multi-site acquisition that has continued well beyond the original protocol description. Because uniformity could not be verified, configuration was carried forward as a measured attribute of each recording and modeled explicitly in all analyses (Sections 2.4 and 2.7).

The protocol documentation describes a 5-point calibration administered before each paradigm and repeated until spatial error fell below 2° at any single point and below 1° on average; calibration outcomes are not reported here and were not used as an inclusion criterion.

### 2.4. Eye-Tracking and Pupillometry Preprocessing

Raw data were preprocessed via a custom Python pipeline following published pupillometry guidelines (Kret & Sjak-Shie, 2019; Mathôt et al., 2018), with one documented departure described below.

Timestamps were anchored to the stimulus onset trigger, and the analysis window was fixed per video at the first 200.0 s (TP), 170.0 s (DM), 116.7 s (WK) and 160.0 s (FF) after onset. Recordings lacking an onset trigger, a recognized sampling rate, or calibration-area metadata were excluded. Gaze and pupil validity were determined independently: zero or missing gaze coordinates, and coordinates falling more than 5% beyond the calibrated screen area, were marked invalid; zero or missing pupil values were marked as blinks. Where a calibrated pupil channel existed, detected blinks were additionally propagated to gaze. Missing-data runs were padded by 100 ms for pupil and 50 ms for gaze to remove partial-closure artifacts. Padding was applied to gaps only, not to outlier flags, since an isolated outlying sample carries no implication about the validity of its neighbors.

Pupil artifacts were then identified in the padded series. A trend-deviation filter flagged samples deviating from a 1 s centered rolling median by more than 16 scaled MADs (Kret & Sjak-Shie, 2019). The dilation-speed filter was implemented as an absolute physiological ceiling of 10 mm/s rather than the MAD-scaled threshold those authors propose. We depart from the published criterion because this tracker reports pupil diameter with sample-and-hold: 33% of consecutive samples at 30 Hz and 41% at 60 Hz are bit-identical, so a large share of inter-sample speeds are exactly zero and the remainder cluster on one or two quantization steps. The MAD then estimates the quantization step rather than physiological variability, and the resulting threshold rejects ordinary movement, 19% of samples in this dataset, at rates differing by sampling rate and therefore confounded with tracker configuration. An absolute ceiling is invariant to both sampling rate and quantization, rejects 0.00% of samples at either rate in these data, and is meaningful here precisely because the pupil channel is restricted to CR-corrected recordings reporting calibrated millimeters. As a safeguard, either filter was skipped for a recording in which it would have rejected more than 20% of samples.

Gaps were linearly interpolated only where the entire run was ≤500 ms; longer gaps were left unfilled rather than partially filled from each end, and leading and trailing runs were never extrapolated. All channels were then low-pass filtered at 10 Hz (second-order zero-phase Butterworth) at the native sampling rate, equalizing effective bandwidth across the three sampling rates, and resampled to a common 30 Hz grid by timestamp-anchored boxcar averaging. Resampling by timestamp rather than by sample index ensures that dropped samples produce empty bins rather than shifting the entire subsequent timeline relative to the stimulus.

Gaze position was expressed in normalized screen coordinates. Because the complex correlation described below is invariant to isotropic but not anisotropic rescaling, and calibration aspect ratio varies across recordings, raw pixel coordinates would perturb both the magnitude and the phase of the resulting statistic. Degrees of visual angle, computed from each recording’s own calibration area, physical stimulus dimensions and head distance, were retained for the coarse oculomotor measures.

Recordings with >10% missing gaze on the output grid were excluded. The same threshold was applied separately to pupil; failing it removed the pupil channel only, leaving gaze data intact. Among CR-corrected recordings, detected blinks accounted for 4.55% of raw pupil samples, gap padding for a further 6.09%, the trend filter for 0.10%, and the speed filter for 0.00%; after interpolation of short gaps, median pupil missingness on the output grid was 1.9% (mean 3.4%). A participant flow chart with exclusion reasons at each stage is provided in Supplementary Figure S1.

### 2.5. Inter-Subject Correlation (ISC) Analysis

To quantify how closely an individual’s gaze and pupillary responses aligned with normative responses, we employed an ISC framework, which extracts a stimulus-driven normative reference trajectory against which individual entrainment is measured.

Every participant, TD or clinical, was scored against a reference constructed from the same number of contributing individuals: for a TD participant, the mean of the remaining *n* − 1 TD participants (strict leave-one-out, preventing circularity); for a clinical participant, the mean of *n* − 1 TD participants, with the omitted participant determined by deterministic rotation so that the procedure is reproducible and the omission is distributed evenly across the TD group. Correlations were computed over the time points at which both the participant and the reference were valid.

Pupil diameter time series were within-subject Z-scored to account for baseline differences in pupil size and luminance reactivity, and pupil ISC was computed as a Pearson correlation between the individual’s Z-scored series and the normative trajectory. The pupil reference was built exclusively from TD participants with a usable pupil channel (CR-corrected and passing the pupil quality threshold of Section 2.4), using the same leave-one-out and rotation scheme.

Traditional gaze ISC computes independent correlations for the horizontal and vertical axes, which does not capture coupled spatiotemporal dynamics. We therefore implemented a two-dimensional complex correlation. At each time point *t*, horizontal and vertical coordinates were mapped into a single complex value *z*(*t*) = *x*(*t*) + *i y*(*t*). With *z*₁ the individual’s complex gaze trajectory and *z*₂ the normative trajectory, both mean-centered, the complex correlation coefficient was

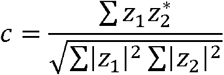

where * denotes the complex conjugate. The modulus |*c*| (range 0–1) quantifies the overall strength of spatiotemporal entrainment: high values indicate that the participant reliably tracked the same screen locations as the normative group. The argument of *c* is the rotation in the screen plane that best aligns the normative trajectory with the participant’s own; we denote its absolute value |θ| and refer to it as absolute gaze phase.

Because the argument of a complex correlation is estimated with precision that depends on its modulus, |θ| and |*c*| are not independent, and gaze position is strongly autocorrelated (median lag-1 autocorrelation 0.976–0.985 across the four videos), which further reduces the effective degrees of freedom available to the phase estimate. We therefore evaluated |θ| against a magnitude-matched null rather than interpreting it directly. For each video, we simulated surrogate participants as first-order autoregressive processes whose gaze related to the reference with a given modulus and the measured autocorrelation but with no true angular offset, across a grid of moduli spanning the 0.5th to 99.5th percentile of the observed values, and interpolated the expected |θ| for each participant’s own |*c*|. We additionally residualized |θ| on a cubic spline in |*c*| (five degrees of freedom) within each video, which requires no simulation assumption. Group models were fitted to the observed |θ|, to the difference from the simulated floor, and to the spline residual.

### 2.6. Coarse Oculomotor Measures

To establish what the inter-subject framework adds over stimulus-independent summaries of eye movement, we computed ten non-inter-subject measures from the same 30 Hz output grid: rate of resolvable gaze shifts (speed exceeding 30 and 60 °/s, counted as runs so that one movement counts once), mean dwell duration between shifts, proportion of time shifting, mean scan-path speed, bivariate contour ellipse area (BCEA, 68%), horizontal and vertical positional standard deviation, overall positional dispersion, and proportion of time within the central third of the screen in both dimensions. Dispersion and central bias were computed in normalized screen coordinates and the remaining measures in degrees of visual angle. Mean dwell duration was defined as valid viewing time divided by the number of shifts and is therefore the exact per-subject reciprocal of gaze-shift rate. Because these were computed at 30 Hz following a common 10 Hz low-pass, they index resolvable gaze shifts rather than individual saccades, and dwell segments rather than fixations in the oculomotor sense; absolute values are therefore not comparable with published norms. They are, however, comparable across participants, since every recording passed through an identical pipeline at an identical final sampling rate, which is what the group comparison requires.

These measures also permit a directional test of one account of the interaction, under which more frequent gaze shifting in ADHD interrupts prolonged idiosyncratic fixation in autism. That account predicts more gaze shifting in the comorbid group than in the autism-without-ADHD group, and we report the standardized difference between those two groups on each measure.

### 2.7. Statistical Analysis

All analyses were conducted in Python using statsmodels. Models were computed separately for each video to preserve the context-specific nature of the paradigms, with missing data handled by listwise deletion applied per metric, so that participants lacking a pupil channel retained their gaze data. Dependent variables and continuous trait scores were standardized within video, so regression coefficients are interpretable in standard-deviation units of the dependent variable. All models adjusted for participant age, biological sex, tracker configuration (the four rate-by-CR combinations present) and screen shape (4:3 versus wider). The two acquisition terms were entered additively rather than crossed, because their crossing yields a cell of 50 recordings cohort-wide. Pupil models additionally adjusted for the proportion of missing pupil samples. Benjamini–Hochberg false discovery rate corrections were applied independently for each model term within each metric family across the four viewing contexts.

Effect sizes are reported both as standardized coefficients and as partial eta-squared. The two diverge in this design: the autism-without-ADHD group comprises 45–78 participants per video against totals of 854–1455, so a large standardized group difference accounts for a small proportion of total variance. Both quantities are reported throughout and neither alone should be read as the effect size. Ninety-five percent confidence intervals for standardized coefficients are Wald intervals from the fitted models.

Two preliminary analyses characterized the sample. Per-video logistic regressions tested whether retention of a recording through preprocessing depended on SRS, SWAN, age or sex, and chi-square tests with Cramér’s V assessed whether acquisition variables were balanced across diagnostic groups.

#### 2.7.1. Dimensional Analysis

To evaluate continuous, transdiagnostic relationships between trait severity and synchrony, we fitted multiple linear regression models in three tiers. Separate models entered SRS and SWAN independently, to assess the broad association of each trait domain. Joint models entered both simultaneously, forcing the traits to compete for variance and isolating effects unique to each domain. Interaction models added an SRS × SWAN term to the joint models, testing whether the two trait domains combine non-additively. Dimensional analyses included the full cohort, including participants in the “Other” diagnostic category, to capture the broadest range of transdiagnostic phenotypic variance; because the categorical models exclude that group, we repeated the dimensional models without it so that the two frameworks could be compared on a common sample.

#### 2.7.2. Categorical Analysis

To relate the dimensional findings to clinical diagnoses, we conducted a 2 × 2 factorial analysis of covariance with participants in the heterogeneous “Other” category excluded. The remaining sample was stratified by the presence or absence of an autism diagnosis (Factor 1) and an ADHD diagnosis (Factor 2), each entered as a 0/1 indicator together with their product. Under this coding, the autism coefficient is the difference between the autism-without-ADHD group and TD controls, the ADHD coefficient is the difference between the ADHD-without-autism group and TD controls, and the interaction coefficient is the amount by which the comorbid group departs from the sum of the two; we refer to the first two as main effects for brevity. A positive interaction accompanying a negative autism coefficient therefore indicates sub-additivity.

Five sensitivity analyses were conducted. First, because tracker configuration was unbalanced across diagnostic groups and the TD reference could not be configuration-matched (some strata contain fewer than ten TD participants per video), models were repeated within the single dominant configuration and, conversely, in the full sample with the acquisition covariates removed. Second, models were repeated with oppositional/conduct diagnosis and CBCL Total T-score as additional covariates; these are reported as sensitivity rather than primary analyses, since CBCL partly indexes the conditions being contrasted. Third, models were repeated with same-day stimulant status as a three-level covariate (stimulant, no stimulant, unknown). Fourth, models were refitted by Huber robust regression to down-weight large residuals, given the size of the autism-without-ADHD cell. Fifth, the autism-without-ADHD and comorbid groups were compared on SRS subscales, full-scale IQ, CBCL total, anxiety and oppositional/conduct diagnoses, and total diagnosis count, using Welch’s *t*-tests with Mann–Whitney tests alongside, to assess whether severity or ascertainment differences could account for any interaction.

## 3. Results

### 3.1. Sample Characteristics and Data Retention

The analytic sample consisted of 2476 participants (1565 male, 911 female) with a mean age of 10.39 years (SD = 3.44). Continuous trait measures reflected a broad phenotypic spectrum, with a mean SRS Total T-score of 57.22 (SD = 11.33) for autistic traits and a mean SWAN Total score of 0.48 (SD = 1.00) for ADHD traits.

Diagnostic groups comprised typically developing controls (n = 239, 9.7%), ADHD without ASD (n = 1156, 46.7%), ASD without ADHD (n = 99, 4.0%), comorbid ASD+ADHD (n = 267, 10.8%), and psychiatric “Other” (n = 715, 28.9%). The ASD-without-ADHD group is the smallest cell by a wide margin, consistent with the high rate of attentional symptoms in autistic children and with the referral structure of a help-seeking cohort; its size is the principal constraint on the precision of the categorical analyses and is revisited in the Limitations.

The volume of viable eye-tracking data varied across the four viewing conditions. Of recordings identified, 77.6% (TP), 72.4% (DM), 75.3% (WK) and 54.5% (FF) survived preprocessing; the markedly lower yield for *Fun with Fractals* reflects exclusion rather than availability, since a comparable number of recordings was identified for that paradigm (n = 2617) as for *The Present* (n = 2729). Requiring clinical characterization and complete covariates gave analytic gaze samples of 1775 (TP), 1959 (DM), 2042 (WK) and 1186 (FF). Between 65% and 69% of these recordings carried corneal-reflection correction (55% for *Fun with Fractals*), reflecting the absence of that stage in one tracker configuration; of those, 84–94% yielded a usable pupil channel, giving pupil samples of 1083, 1176, 1308 and 545. Excluding the heterogeneous “Other” group leaves categorical subsamples of 1257, 1390, 1455 and 854 respectively. Demographic and clinical distributions across these subsets are given in Table 1, and a participant flow chart with exclusion reasons at each stage in Supplementary Figure S1.

**Table 1.**
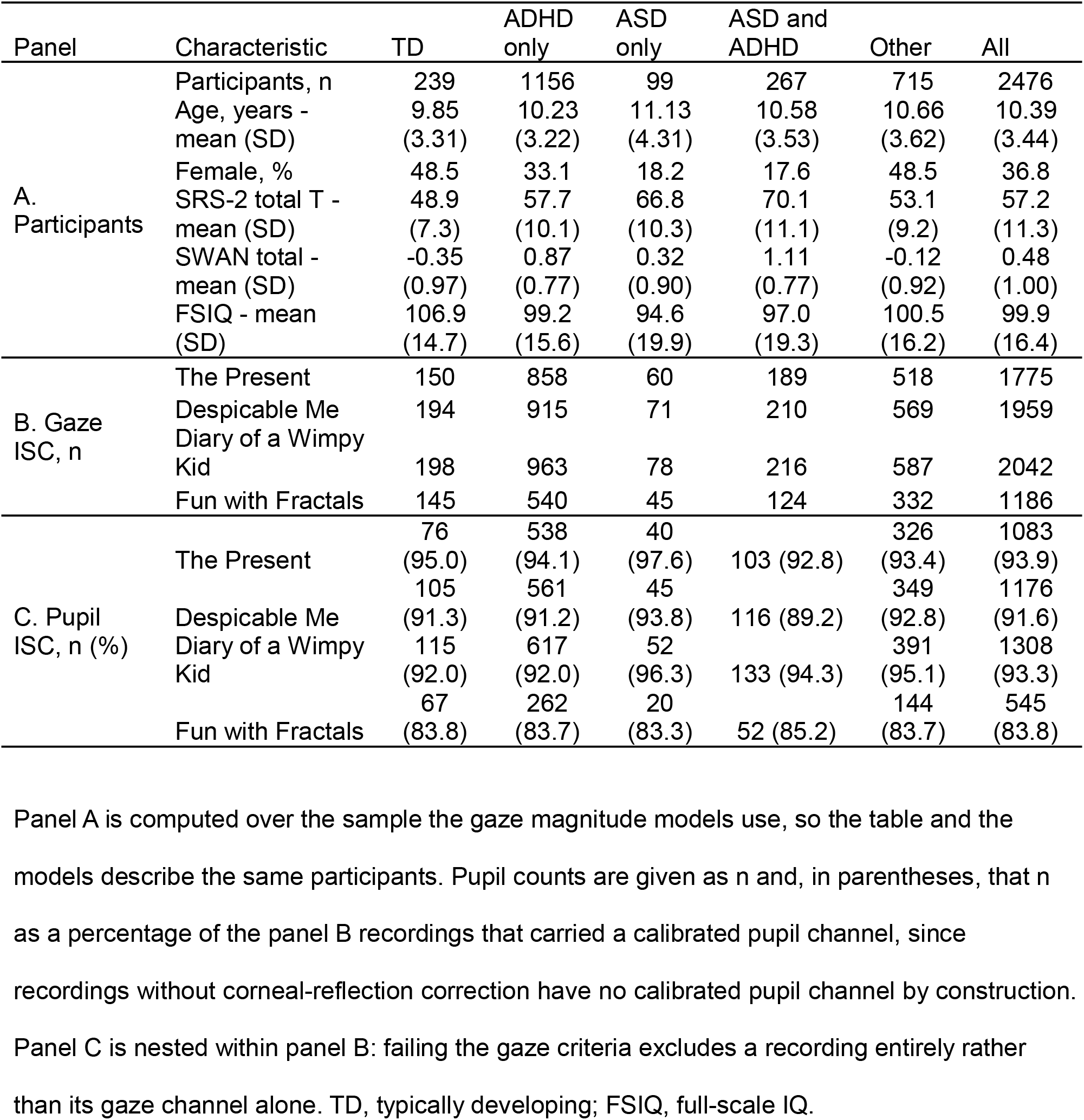
Characteristics of the analysis sample and number of recordings by diagnostic group.

| Panel | Characteristic | TD | ADHD only | ASD only | ASD and ADHD | Other | All |
|---|---|---|---|---|---|---|---|
| A. Participants | Participants, n | 239 | 1156 | 99 | 267 | 715 | 2476 |
|  | Age, years - mean (SD) | 9.85 (3.31) | 10.23 (3.22) | 11.13 (4.31) | 10.58 (3.53) | 10.66 (3.62) | 10.39 (3.44) |
|  | Female, % | 48.5 | 33.1 | 18.2 | 17.6 | 48.5 | 36.8 |
|  | SRS-2 total T - mean (SD) | 48.9 (7.3) | 57.7 (10.1) | 66.8 (10.3) | 70.1 (11.1) | 53.1 (9.2) | 57.2 (11.3) |
|  | SWAN total - mean (SD) | -0.35 (0.97) | 0.87 (0.77) | 0.32 (0.90) | 1.11 (0.77) | -0.12 (0.92) | 0.48 (1.00) |
|  | FSIQ - mean (SD) | 106.9 (14.7) | 99.2 (15.6) | 94.6 (19.9) | 97.0 (19.3) | 100.5 (16.2) | 99.9 (16.4) |
| B. Gaze ISC, n | The Present | 150 | 858 | 60 | 189 | 518 | 1775 |
|  | Despicable Me | 194 | 915 | 71 | 210 | 569 | 1959 |
|  | Diary of a Wimpy Kid | 198 | 963 | 78 | 216 | 587 | 2042 |
|  | Fun with Fractals | 145 | 540 | 45 | 124 | 332 | 1186 |
| C. Pupil ISC, n (%) | The Present | 76 (95.0) | 538 (94.1) | 40 (97.6) | 103 (92.8) | 326 (93.4) | 1083 (93.9) |
|  | Despicable Me | 105 (91.3) | 561 (91.2) | 45 (93.8) | 116 (89.2) | 349 (92.8) | 1176 (91.6) |
|  | Diary of a Wimpy Kid | 115 (92.0) | 617 (92.0) | 52 (96.3) | 133 (94.3) | 391 (95.1) | 1308 (93.3) |
|  | Fun with Fractals | 67 (83.8) | 262 (83.7) | 20 (83.3) | 52 (85.2) | 144 (83.7) | 545 (83.8) |

Panel A is computed over the sample the gaze magnitude models use, so the table and the models describe the same participants. Pupil counts are given as n and, in parentheses, that n as a percentage of the panel B recordings that carried a calibrated pupil channel, since recordings without corneal-reflection correction have no calibrated pupil channel by construction. Panel C is nested within panel B: failing the gaze criteria excludes a recording entirely rather than its gaze channel alone. TD, typically developing; FSIQ, full-scale IQ.

Data availability was not independent of phenotype. In per-video logistic regressions of retention on SRS, SWAN, age and sex, higher autistic traits predicted lower retention in three of four videos (DM: b = −0.011, p = 0.013; WK: b = −0.016, p = 0.0003; FF: b = −0.015, p = 0.0007), with no significant association for TP (p = 0.16) and no SWAN effect in any video (all p > 0.11; Supplementary Table S1).

### 3.2. Naturalistic Viewing Entrains Robust Spatiotemporal and Pupillary Synchrony

To verify that naturalistic viewing elicited shared physiological states, we first evaluated baseline inter-subject synchrony. Visual inspection of the raw trajectories indicated robust entrainment: as illustrated in Figure 1 for *The Present*, vertical banding is clearly visible in the subject-by-time matrices for both gaze and pupil, marking time points at which many participants moved or dilated together. Participants are sorted by autistic trait severity, shown alongside the matrices, so that the ordering can be inspected directly rather than assumed.

**Figure 1.**
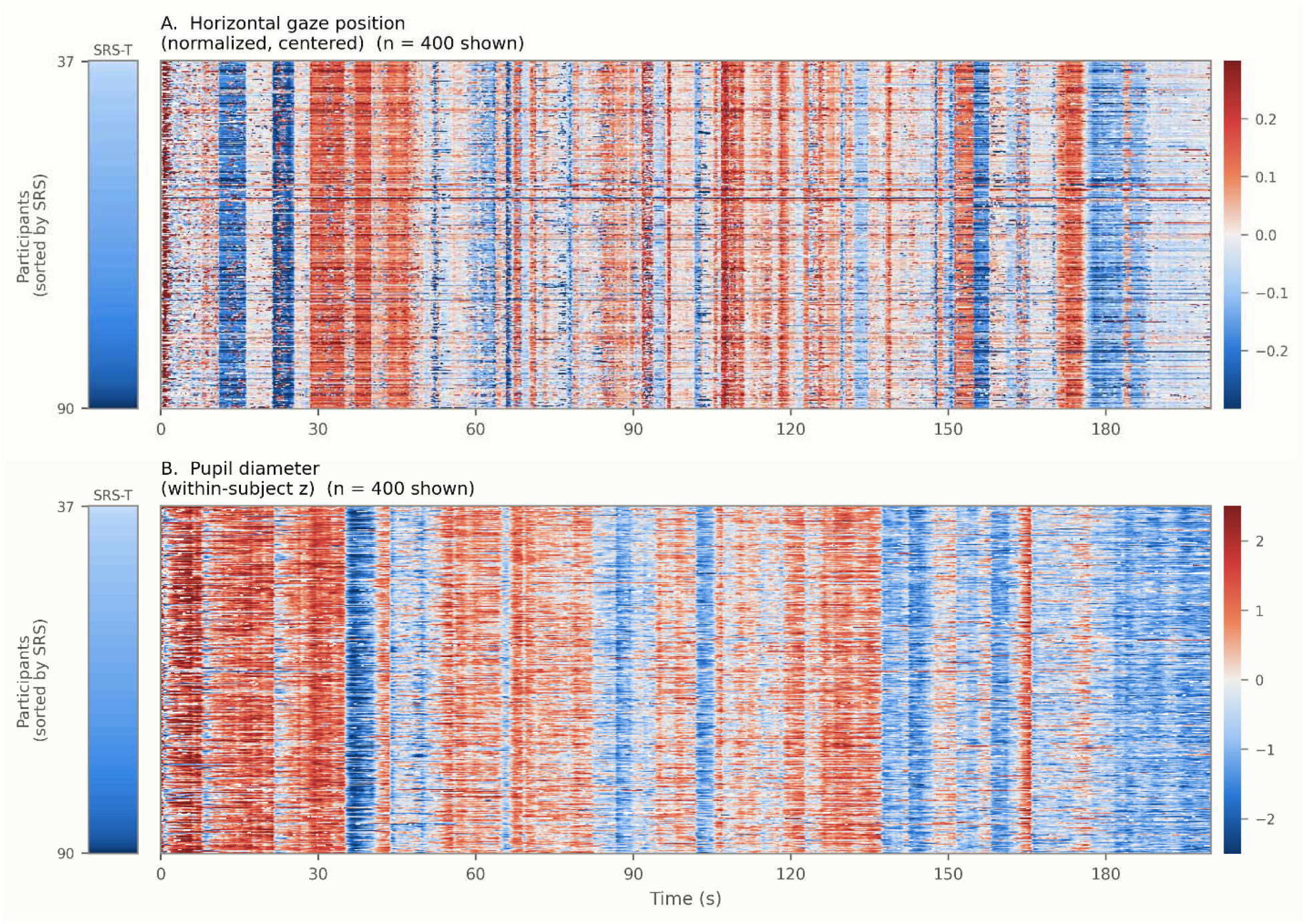
Gaze and pupil dynamics during naturalistic viewing of *The Present*. (**A**) Horizontal gaze position and (**B**) pupil diameter, as subject-by-time matrices. Each row is one participant’s time series on the common 30 Hz grid; time runs left to right in seconds from stimulus onset. Rows are ordered by Social Responsiveness Scale (SRS-2) Total T-score, lowest at top, and the sort variable itself is drawn as the narrow strip at the left of each panel so the ordering can be inspected rather than assumed. Color encodes signed deviation: gaze from the group median position (±0.3 in normalized screen units), pupil from each participant’s own mean (±2.5 within-subject SD). Vertical banding marks time points at which many participants moved or dilated together, which is the entrainment the inter-subject analyses quantify. Panel B is restricted to recordings with a corneal-reflection corrected pupil channel, since recordings without that correction report pupil size in camera pixels rather than a calibrated diameter. For legibility, 400 participants are drawn per panel, selected at evenly spaced positions in the SRS ordering of all participants with an SRS-2 score, so that the distribution of scores is preserved; the selection is deterministic.

Gaze ISC magnitude demonstrated substantial synchrony across the sample in every context, and varied across the four videos. Entrainment was highest during the emotionally complex animated short *The Present* (M = 0.664, SD = 0.079), intermediate and closely matched during *Despicable Me* (M = 0.601, SD = 0.087) and *Diary of a Wimpy Kid* (M = 0.602, SD = 0.084), and lowest during the abstract, non-social *Fun with Fractals* (M = 0.505, SD = 0.082).

This ordering is not attributable to differences in reference reliability. The typically developing reference comprised a comparable number of participants for *The Present* (n = 150) and *Fun with Fractals* (n = 145; Table 1), yet mean synchrony differed by 0.16 between them, whereas *Despicable Me* and *Diary of a Wimpy Kid* were built on the two largest references (n = 194 and 198) and yielded intermediate values. Reference size and synchrony level are therefore unrelated across the four contexts, and the gradient is consistent with the social and narrative content of the stimulus rather than with the precision of the normative trajectory.

Absolute gaze phase was low on average but highly dispersed, with standard deviations exceeding means in every context (TP: M = 0.085 rad, SD = 0.146; DM: M = 0.092, SD = 0.167; WK: M = 0.086, SD = 0.132; FF: M = 0.106, SD = 0.161). We defer interpretation of this metric to Section 3.5, where it is evaluated against the phase expected from synchrony magnitude alone.

Pupil ISC was strongly positive throughout. Pupillary entrainment was highest during the two social narratives, *The Present* (M = 0.757, SD = 0.103) and *Diary of a Wimpy Kid* (M = 0.779, SD = 0.100), and lower during the fast-paced *Despicable Me* (M = 0.557, SD = 0.139) and the abstract *Fun with Fractals* (M = 0.620, SD = 0.119).

Having established that the stimuli drove synchrony in both channels and did so differentially, we next examined how these metrics varied with autistic and ADHD traits.

### 3.3. Dimensional Analysis of SRS and SWAN

Continuous trait models were fitted with each domain entered alone and with both entered simultaneously, adjusted for age, sex, tracker configuration and screen shape. Figure 2 summarizes the standardized coefficients for every metric and context under both specifications; full statistics are given in Supplementary Tables S3 and S4, and the underlying scatter plots in Supplementary Figures S2 and S3. These models retain participants in the “Other” diagnostic category and require complete SRS and SWAN reports, so their samples are slightly smaller than the gaze samples in Table 1 (n = 1144–1971 against 1186–2042).

**Figure 2.**
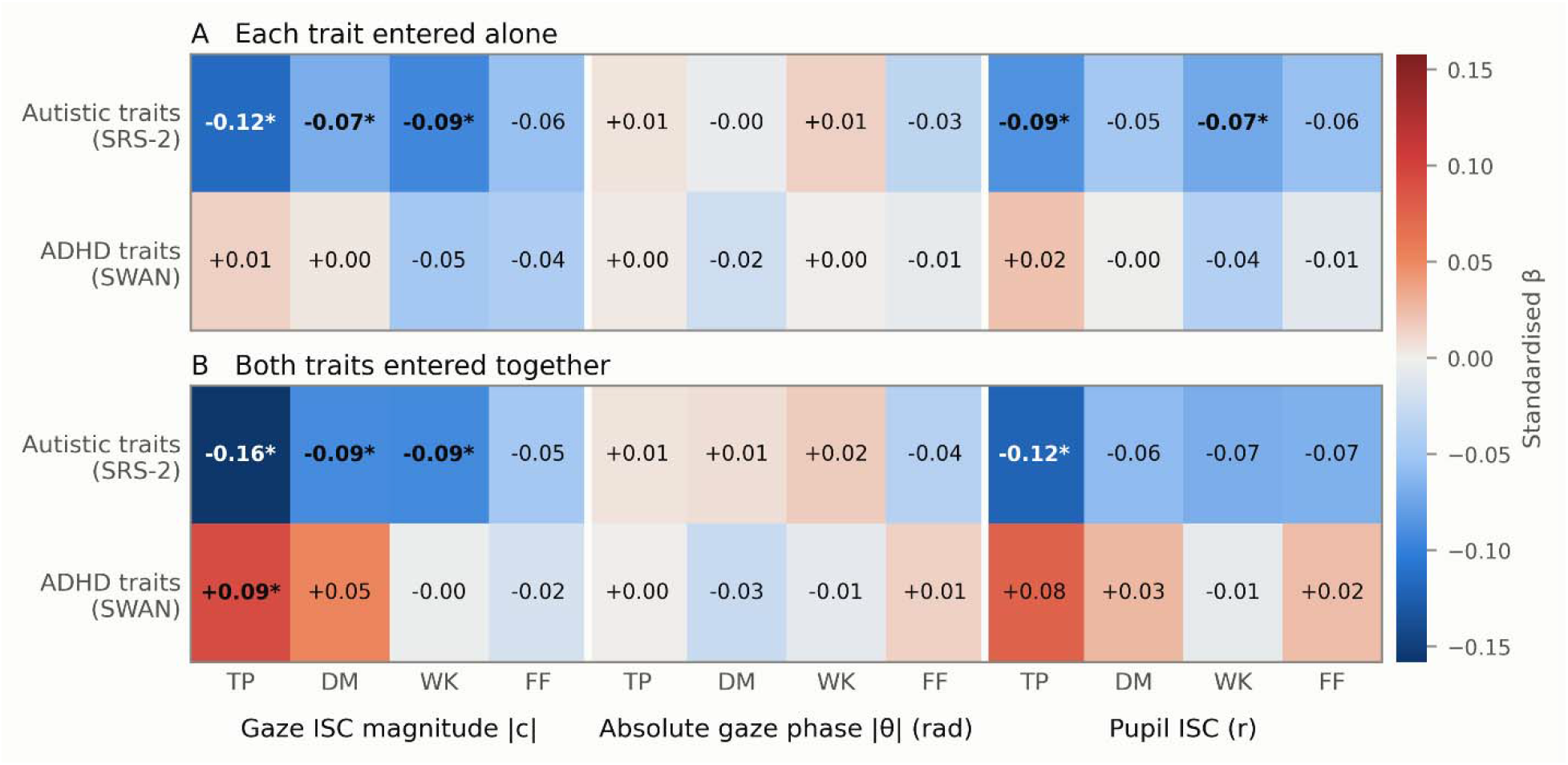
Dimensional associations between neurodevelopmental traits and synchrony, across all metrics and viewing paradigms. Cells are standardized regression coefficients to two decimal places, adjusted for age, sex, tracker configuration and screen shape; asterisks and bold mark associations surviving Benjamini-Hochberg correction within each metric family across the four paradigms. (**A**) Each trait domain entered in its own model. (**B**) Both entered simultaneously, so each coefficient is the association net of the other. Reading a column from A to B shows the consequence of the r = 0.46–0.47 correlation between the trait scores: autistic-trait coefficients strengthen while ADHD coefficients reverse sign. Paradigms are *The Present* (TP), *Despicable Me* (DM), *Diary of a Wimpy Kid* (WK) and *Fun with Fractals* (FF). Exact coefficients and full statistics are in Supplementary Tables S3 and S4; the underlying scatter plots are in Supplementary Figures S2 and S3.

Entered alone, autistic traits predicted reduced gaze synchrony in three of the four contexts (Figure 2A): *The Present* (β = −0.116, q = 5.2 × 10⁻ ), *Despicable Me* (β = −0.069, q = 0.004) and *Diary of a Wimpy Kid* (β = −0.095, q = 4.1 × 10⁻ ), with the same sign but not significant during *Fun with Fractals* (β = −0.055, q = 0.061). ADHD traits entered alone showed no association in any context (|β| ≤ 0.048, all q ≥ 0.14).

Entering both domains simultaneously sharpened the autistic-trait association rather than diluting it, and reversed the sign of the ADHD coefficient (Figure 2B). In *The Present* the SRS coefficient increased in magnitude from −0.116 to −0.158 (q = 2.3 × 10⁻ ) while the SWAN coefficient moved from +0.015 (q = 0.74) to +0.092 (q = 0.004): with autistic traits held constant, higher ADHD traits predicted *higher* gaze synchrony. The same pattern appeared in *Despicable Me* without reaching significance for the ADHD term (SRS −0.069 to −0.092, q = 0.0006; SWAN +0.004 to +0.051, q = 0.13), and was absent in the remaining two contexts. This is the signature of statistical suppression between two positively correlated predictors, which the SRS and SWAN totals are here (*r* = 0.46–0.47), and it is the continuous counterpart of the categorical interaction reported in Section 3.4.

Pupillary synchrony showed the same direction over a narrower range. Autistic traits entered alone predicted reduced pupil ISC during *The Present* (β = −0.087, q = 0.017) and *Diary of a Wimpy Kid* (β = −0.072, q = 0.022), but not during *Despicable Me* or *Fun with Fractals* (both q ≥ 0.14). The joint models again strengthened the autistic-trait coefficient in *The Present* (β = −0.120, q = 0.002), with a positive but non-significant ADHD term (β = +0.076, q = 0.11); the association in *Diary of a Wimpy Kid* fell short of significance once ADHD traits were entered (β = −0.069, q = 0.058).

Absolute gaze phase showed no dimensional association with either trait domain, in any context, under either specification (all |β| ≤ 0.038, all q ≥ 0.86). Its status is addressed in Section 3.5.

Adding an SRS × SWAN product term to the joint models yielded no significant interaction for any metric in any context (all |β| ≤ 0.045, all q ≥ 0.21; Supplementary Table S5).

Because the categorical models in Section 3.4 exclude the “Other” group, we repeated the dimensional models without it (Supplementary Table S6). The gaze results were unchanged: autistic traits remained significant in the same three contexts, and the positive ADHD-trait coefficient in the joint model for *The Present* strengthened (β = +0.117, q = 0.001). The pupil associations did not survive the restriction (single-trait models, all q ≥ 0.18; joint model for *The Present*, q = 0.064) and should be regarded as holding in the full referred sample only.

### 3.4. Categorical Analysis of ASD and ADHD

A 2 × 2 analysis of covariance contrasted typically developing controls, autism without ADHD, ADHD without autism, and the comorbid group, adjusted for age, sex, tracker configuration and screen shape, with the heterogeneous “Other” group excluded (n = 854–1455; Table 2, Figure 3).

**Figure 3.**
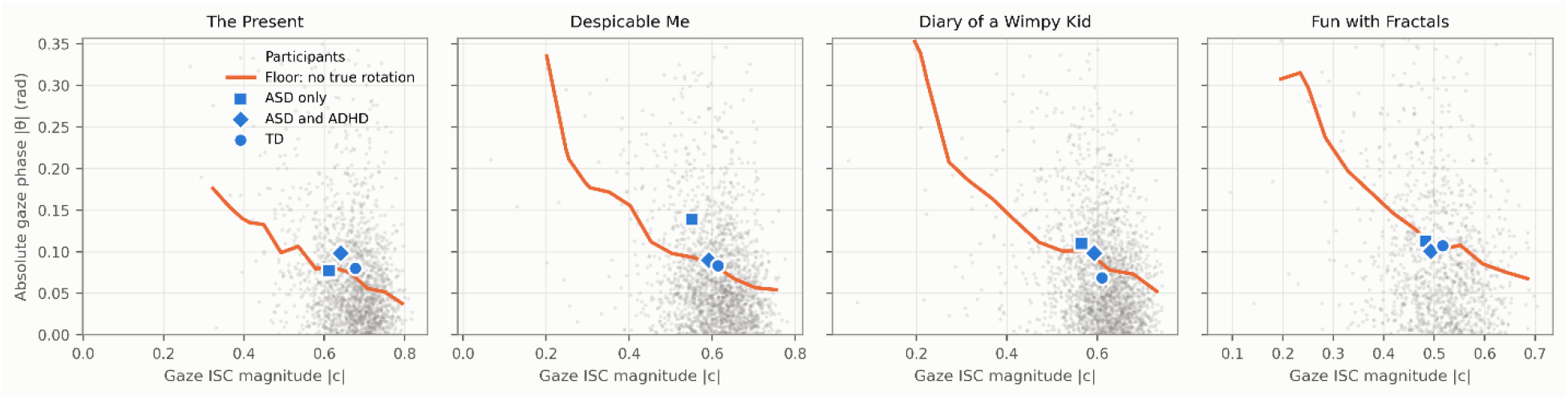
Gaze and pupil synchrony by ASD and ADHD status. Rows: gaze ISC magnitude |*c*| (top) and pupil ISC (bottom). Columns: the four viewing contexts. Within each panel, autism status is encoded by color and ADHD status by horizontal position, so a sub-additive interaction appears as a narrower vertical gap between the two colors on the right of the panel than on the left. Points are individual participants; boxes give the median and interquartile range; markers and vertical bars give the cell mean and its 95% confidence interval. Point opacity, size and jitter width are scaled by cell size, because the four cells differ by an order of magnitude (45–78 against 540–963) and a fixed opacity would render the largest cell as a solid block and the smallest as a few specks. Cell *n* is printed along the bottom of each panel. Axis limits span the 0.5th to 99.5th percentile of the pooled distribution. The heterogeneous “Other” diagnostic group is excluded. Models adjusted for age, sex, tracker configuration and screen shape; see Table 2 for estimates.

**Table 2.** Two-by-two (ASD × ADHD) analysis of covariance for each synchrony metric.

| Metric | Term | Video | n | β | 95% CI | η<sub>p</sub><sup>2</sup> | p | q |
|---|---|---|---|---|---|---|---|---|
| Gaze ISC Magnitude | ASD | TP | 1257 | -0.784 | [-1.08, -0.49] | 0.021 | <0.001 | <0.001 |
|  | ASD | DM | 1390 | -0.785 | [-1.06, -0.51] | 0.022 | <0.001 | <0.001 |
|  | ASD | WK | 1455 | -0.602 | [-0.86, -0.34] | 0.014 | <0.001 | <0.001 |
|  | ASD | FF | 854 | -0.451 | [-0.79, -0.12] | 0.008 | 0.009 | 0.009 |
|  | ADHD | TP | 1257 | -0.121 | [-0.29, 0.05] | 0.002 | 0.174 | 0.232 |
|  | ADHD | DM | 1390 | -0.160 | [-0.32, -0.00] | 0.003 | 0.048 | 0.191 |
|  | ADHD | WK | 1455 | -0.076 | [-0.23, 0.08] | 0.001 | 0.332 | 0.332 |
|  | ADHD | FF | 854 | -0.133 | [-0.32, 0.05] | 0.002 | 0.162 | 0.232 |
|  | ASD × ADHD | TP | 1257 | 0.497 | [0.16, 0.83] | 0.007 | 0.004 | 0.005 |
|  | ASD × ADHD | DM | 1390 | 0.649 | [0.34, 0.96] | 0.012 | <0.001 | <0.001 |
|  | ASD × ADHD | WK | 1455 | 0.459 | [0.16, 0.75] | 0.006 | 0.002 | 0.005 |
|  | ASD × ADHD | FF | 854 | 0.277 | [-0.11, 0.66] | 0.002 | 0.159 | 0.159 |
| Absolute Gaze Phase | ASD | TP | 1257 | -0.005 | [-0.31, 0.30] | 0.000 | 0.976 | 0.976 |
|  | ASD | DM | 1390 | 0.406 | [0.13, 0.68] | 0.006 | 0.004 | 0.016 |
|  | ASD | WK | 1455 | 0.351 | [0.09, 0.62] | 0.005 | 0.009 | 0.019 |
|  | ASD | FF | 854 | 0.066 | [-0.27, 0.41] | 0.000 | 0.704 | 0.938 |
|  | ADHD | TP | 1257 | 0.065 | [-0.11, 0.24] | 0.000 | 0.474 | 0.634 |
|  | ADHD | DM | 1390 | 0.058 | [-0.10, 0.22] | 0.000 | 0.476 | 0.634 |
|  | ADHD | WK | 1455 | 0.137 | [-0.02, 0.29] | 0.002 | 0.086 | 0.345 |
|  | ADHD | FF | 854 | -0.015 | [-0.20, 0.17] | 0.000 | 0.873 | 0.873 |
|  | ASD × ADHD | TP | 1257 | 0.040 | [-0.30, 0.38] | 0.000 | 0.816 | 0.816 |
|  | ASD × ADHD | DM | 1390 | -0.387 | [-0.70, -0.08] | 0.004 | 0.015 | 0.060 |
|  | ASD × ADHD | WK | 1455 | -0.245 | [-0.55, 0.06] | 0.002 | 0.111 | 0.222 |
|  | ASD × ADHD | FF | 854 | -0.091 | [-0.48, 0.30] | 0.000 | 0.648 | 0.816 |
| Pupil ISC | ASD | TP | 757 | -0.814 | [-1.19, -0.44] | 0.024 | <0.001 | <0.001 |
|  | ASD | DM | 827 | -0.476 | [-0.82, -0.13] | 0.009 | 0.007 | 0.015 |
|  | ASD | WK | 917 | -0.351 | [-0.68, -0.02] | 0.005 | 0.038 | 0.051 |
|  | ASD | FF | 401 | -0.151 | [-0.66, 0.36] | 0.001 | 0.561 | 0.561 |
|  | ADHD | TP | 757 | -0.079 | [-0.31, 0.16] | 0.001 | 0.511 | 0.982 |
|  | ADHD | DM | 827 | -0.002 | [-0.21, 0.21] | 0.000 | 0.985 | 0.985 |
|  | ADHD | WK | 917 | -0.081 | [-0.28, 0.12] | 0.001 | 0.433 | 0.982 |
|  | ADHD | FF | 401 | 0.047 | [-0.23, 0.32] | 0.000 | 0.737 | 0.982 |
|  | ASD × ADHD | TP | 757 | 0.559 | [0.13, 0.99] | 0.009 | 0.010 | 0.041 |
|  | ASD × ADHD | DM | 827 | 0.102 | [-0.30, 0.50] | 0.000 | 0.615 | 0.789 |
|  | ASD × ADHD | WK | 917 | 0.284 | [-0.10, 0.66] | 0.002 | 0.142 | 0.283 |
|  | ASD × ADHD | FF | 401 | 0.081 | [-0.51, 0.67] | 0.000 | 0.789 | 0.789 |

Primary specification. The ‘Other’ diagnostic group is excluded. Models adjust for age, sex, tracker configuration and screen shape. β, standardized coefficient; CI, confidence interval; ηp², partial eta-squared; q, Benjamini–Hochberg corrected p. TP, *The Present*; DM, *Despicable Me*; WK, *Diary of a Wimpy Kid*; FF, *Fun with Fractals*.

An isolated autism diagnosis was associated with reduced gaze synchrony in all four viewing contexts, with large standardized differences: *The Present* (β = −0.78, q = 5.4 × 10⁻ ), *Despicable Me* (β = −0.78, q = 9.4 × 10⁻ ), *Diary of a Wimpy Kid* (β = −0.60, q = 7.5 × 10⁻ ) and *Fun with Fractals* (β = −0.45, q = 0.009). An isolated ADHD diagnosis produced no significant main effect on gaze synchrony in any context (all q ≥ 0.19).

The interaction terms were positive wherever the main effects were negative, indicating sub-additivity. Significant ASD × ADHD interactions emerged during *The Present* (β = +0.50, q = 0.005), *Despicable Me* (β = +0.65, q < 0.001) and *Diary of a Wimpy Kid* (β = +0.46, q = 0.005), with the same sign but no significance during *Fun with Fractals* (β = +0.28, q = 0.159). Decomposing the estimates for *The Present* illustrates the pattern: relative to typically developing controls, autism without ADHD was associated with a reduction of 0.78 SD, whereas the corresponding reduction for comorbid children relative to children with ADHD alone was 0.29 SD. Rather than compounding, the co-occurrence of ADHD accompanied an attenuation of roughly two-thirds of the difference associated with autism alone.

Effect sizes expressed as variance explained were small throughout (gaze ISC: ηp² = 0.008–0.022 for the autism main effect, 0.002–0.012 for the interaction). The divergence between large standardized differences and small variance explained follows from cell composition: the autism-without-ADHD group comprised 45–78 participants against totals of 854–1455, so a substantial group difference necessarily accounts for a small share of total variance. Both quantities are reported throughout.

Pupillary synchrony showed a partially parallel pattern. An isolated autism diagnosis was associated with reduced pupil ISC during *The Present* (β = −0.81, q < 0.001) and *Despicable Me* (β = −0.48, q = 0.015), but not during *Diary of a Wimpy Kid* (β = −0.35, q = 0.051) or *Fun with Fractals*. A significant positive ASD × ADHD interaction emerged in a single context, *The Present* (β = +0.56, q = 0.041). Given that this is one context of four, we treat the pupil interaction as exploratory and do not interpret it further.

Absolute gaze phase showed significant autism main effects during *Despicable Me* (β = +0.41, q = 0.016) and *Diary of a Wimpy Kid* (β = +0.35, q = 0.019), with interactions in the opposite direction that did not survive correction. Because these estimates mirror the magnitude results with reversed sign, their status is addressed in Section 3.5.

### 3.5. Absolute Gaze Phase Is Not Independent of Synchrony Magnitude

The two components of the complex correlation are mathematically coupled: the argument of *c* is estimated with precision that scales with its modulus. We therefore examined whether absolute gaze phase carried information beyond gaze ISC magnitude before interpreting it.

Within each viewing context, |θ| was substantially and negatively associated with |*c*| (Pearson *r* = −0.32 to −0.45; Supplementary Table S11), with the two metrics sharing 10–20% of their variance. Participants with weaker overall entrainment therefore showed larger apparent angular misalignment as a matter of course, independent of any difference in where they looked. The Spearman coefficients were roughly half the Pearson values (ρ = −0.20 to −0.25), indicating that the association is carried disproportionately by participants at the extremes of the distribution — a property that becomes relevant again below.

To quantify the expected magnitude of this artifact we simulated, for each video, surrogate participants whose gaze related to the normative trajectory with a given modulus and the empirically measured gaze autocorrelation (lag-1 *r* = 0.976–0.985), but with no true angular offset. Observed group means in |θ| were close to, and frequently below, the phase these surrogates produced (Figure 4, Supplementary Table S12). The largest positive departure was the autism-without-ADHD group during *Despicable Me*, observed 0.139 against an expected 0.097; that cell contains 71 participants, and its excess is the source of the larger of the two apparent phase main effects in the primary models. Every other group mean in every context departed from its floor by between −0.020 and +0.022 radians. That several groups fell below their simulated floor indicates the parametric surrogate slightly overestimates the floor in absolute terms, which is why the formal tests below rely on within-video contrasts, where a uniform offset cancels, and on a non-parametric alternative requiring no simulation.

**Figure 4.**
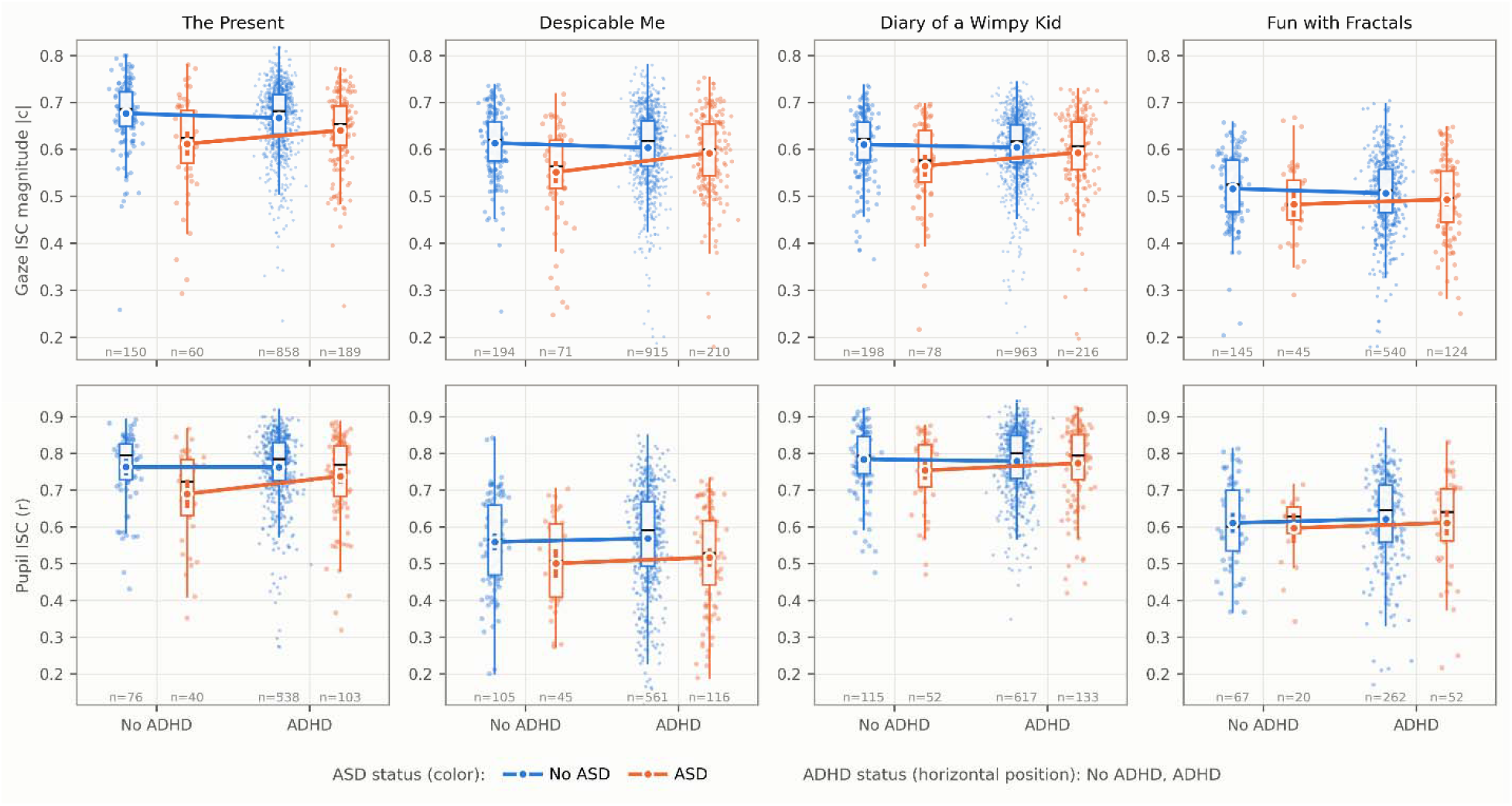
Absolute gaze phase against the floor implied by synchrony magnitude alone. One panel per viewing context. Gray points are individual participants, plotted as absolute phase |θ| against gaze ISC magnitude |*c*|. The orange curve is the phase expected from magnitude alone: for each video, surrogate participants were simulated whose gaze related to the normative trajectory with a given modulus and the empirically measured gaze autocorrelation (lag-1 *r* = 0.976–0.985) but with no true angular offset. Blue markers give group means — square, autism without ADHD; diamond, comorbid; circle, typically developing — positioned at each group’s mean |*c*| and mean |θ|. The group ordering in |θ| follows the group ordering in |*c*|, and the group means lie close to or below the simulated floor, which is why phase is not interpreted as an independent index of spatial misalignment. The vertical axis is truncated at the 97.5th percentile of |θ|. Formal tests used the spline residual of |θ| on |*c*| rather than the simulated floor shown here (Supplementary Tables S13 and S14).

In the primary categorical models, |θ| showed significant autism main effects during *Despicable Me* (β = +0.41, q = 0.016) and *Diary of a Wimpy Kid* (β = +0.35, q = 0.019), in the direction expected if reduced entrainment inflates apparent misalignment. Neither survived adjustment for magnitude. Residualizing |θ| on a cubic spline in |*c*| within each video — which assumes nothing about the data-generating process — eliminated both effects (DM: β = +0.13, q = 0.49; WK: β = +0.15, q = 0.49), and no main effect or interaction on the residual reached significance in any context (all q ≥ 0.14; Supplementary Table S14). Subtracting the simulated floor gave the same result by a different route (all q ≥ 0.064; Supplementary Table S13), with the two approaches disagreeing on which context came closest to significance — the spline favoring *The Present* and the simulation *Despicable Me* — as expected when no stable signal is present.

We therefore do not interpret absolute gaze phase as an independent index of spatial misalignment. The group differences it shows are those its modulus already carries, re-expressed; dimensionally it shows no association with either trait domain at all (Section 3.3); and its apparent effects are four times more sensitive to extreme observations than the magnitude effects computed from the same recordings (Section 3.7). Phase estimates are reported in Table 2 and Supplementary Tables S11–S14 for completeness but are not discussed further.

### 3.6. Coarse Oculomotor Measures

To establish what the inter-subject framework adds over stimulus-independent summaries of eye movement, we applied the same 2 × 2 models to ten non-inter-subject measures derived from the identical 30 Hz time series: gaze-shift rate at two velocity thresholds, mean dwell duration, proportion of time shifting, scan-path speed, BCEA, horizontal and vertical positional standard deviation, overall dispersion, and proportion of time within the central third of the display.

Across 120 tests (ten measures × four contexts × three model terms), two reached significance after correction, both for mean dwell duration during *Diary of a Wimpy Kid*: an autism main effect (β = +0.51, q < 0.001) and an ASD × ADHD interaction (β = −0.47, q = 0.007; Supplementary Table S15). Children with autism and no ADHD held gaze for longer between shifts than typically developing controls, and the co-occurrence of ADHD attenuated that difference. This result requires qualification. Mean dwell duration and gaze-shift rate are exact per-subject reciprocals, and the shift-rate analogues of both terms were null in the same context (autism main effect: β = −0.10, q = 0.96; interaction: β = +0.12, q = 0.71), indicating that the dwell result reflects the upper tail of the dwell distribution rather than a difference in central tendency (Supplementary Table S16). No other measure showed a significant main effect or interaction in any context (all q ≥ 0.11); gaze-shift rate, dispersion, BCEA, and scan-path speed — the measures most directly comparable with standard oculomotor reporting — were undifferentiated by diagnostic group throughout.

We additionally tested the specific prediction that follows from an account in which more frequent gaze shifting disrupts rigid spatial fixation: children with co-occurring autism and ADHD should shift gaze more often than children with autism alone. The direction was consistent in all four contexts (*The Present*: +0.09 shifts/s, d = 0.17; *Despicable Me*: +0.06, d = 0.10; *Diary of a Wimpy Kid*: +0.13, d = 0.29; *Fun with Fractals*: +0.16, d = 0.27; Supplementary Table S16; Supplementary Figure S5), and agreed in direction with the dwell-duration result above. These differences were small, however, and none reached significance in the corresponding 2 × 2 models (shift-rate interaction, all q ≥ 0.46).

Thus the diagnostic structure detected by inter-subject synchrony in the same recordings was not recoverable from coarse summaries of the same gaze data.

### 3.7. Sensitivity Analyses

To establish that the interaction reflected a stable phenotype rather than an artifact of analytic choices, sample composition, or pharmacological status, we conducted four sensitivity analyses and one comparison of clinical characteristics. The interaction estimate under each covariate and sample specification is shown in Figure 5.

**Figure 5.**
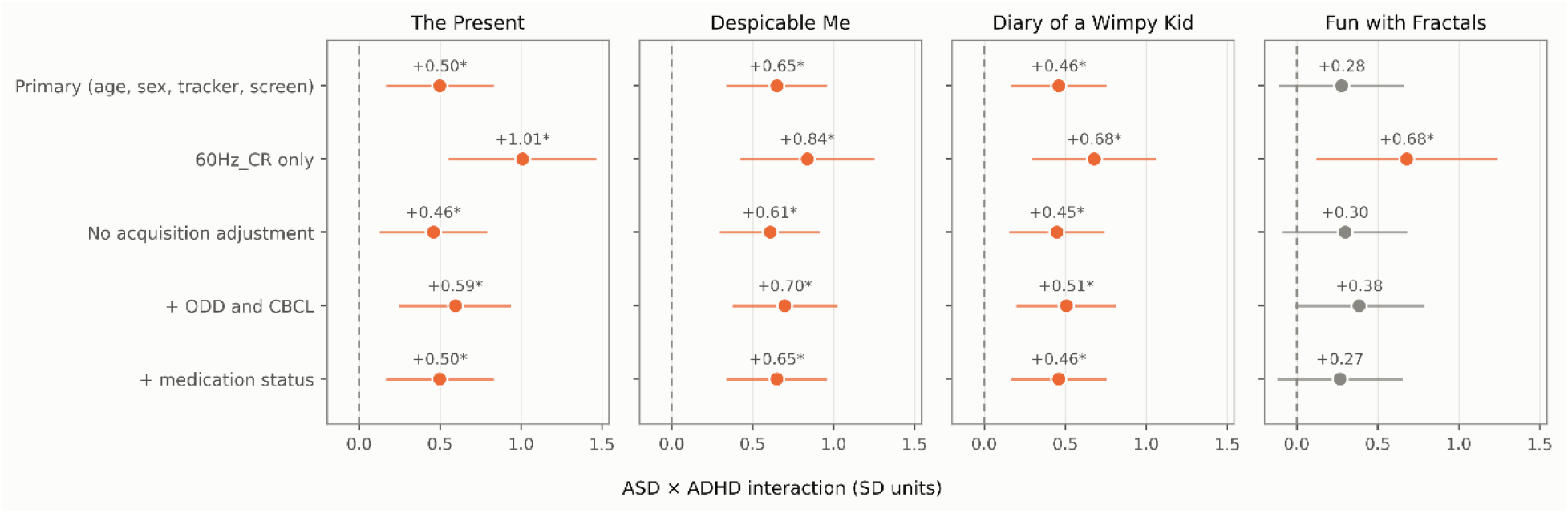
The ASD × ADHD interaction on gaze ISC magnitude across analytic specifications. One panel per viewing context; one row per specification, labeled at the left. Markers are standardized interaction coefficients in standard-deviation units, with the value printed above each; horizontal lines are 95% confidence intervals. Positive values indicate sub-additivity, that is, a smaller reduction associated with autism when ADHD co-occurs. Orange marks estimates significant after Benjamini-Hochberg correction within the metric family (*q* < 0.05, also flagged with an asterisk); gray marks those that are not. The dashed vertical line is the null.

First, because tracker configuration was unbalanced across diagnostic groups and the normative reference could not be configuration-matched, we repeated the models within the single dominant configuration (60 Hz with corneal-reflection correction; Supplementary Table S7). Despite the reduction in sample size (n = 438–820 against 854–1455), estimates increased substantially rather than attenuating. The autism main effect on gaze synchrony ranged from −0.85 to −1.15 SD (all q ≤ 0.001) and the ASD × ADHD interaction from +0.68 to +1.01 SD, reaching significance in all four contexts including *Fun with Fractals* (q = 0.018). Conversely, removing tracker configuration and screen shape from the full-sample models changed the gaze estimates by at most 0.04 SD and altered the significance of no term (Supplementary Table S8). Acquisition heterogeneity therefore attenuated the effects in the full sample rather than generating them.

Second, we repeated the models with oppositional/conduct diagnosis and CBCL Total T-score as additional covariates (Supplementary Table S9). The interaction strengthened in every context (TP: β = +0.59, q = 0.001; DM: β = +0.70, q < 0.001; WK: β = +0.51, q = 0.002; FF: β = +0.38, q = 0.060), as did the autism main effects. General psychopathology therefore suppresses rather than produces the observed pattern. We report this as a sensitivity analysis rather than a primary model because CBCL partly indexes the conditions being contrasted.

Third, we incorporated same-day stimulant status as a three-level covariate (stimulant, no stimulant, unknown; Supplementary Table S20). In the three contexts with a significant interaction, estimates were identical to the primary specification to three decimal places (TP: β = +0.497 against +0.497; DM: +0.649 against +0.649; WK: +0.459 against +0.459); in *Fun with Fractals* the estimate moved by 0.011 SD and remained non-significant. Because status was unknown for 79.2% of the sample, this establishes that stimulant exposure does not account for the interaction among participants whose status is documented; it does not exclude pharmacological effects in the remainder.

Fourth, we refitted the models by Huber robust regression to down-weight extreme observations, given the size of the autism-without-ADHD cell (Supplementary Table S10). Estimates attenuated by 10–20% in most contexts relative to ordinary least squares, as expected when influential tail observations are down-weighted, but the inferential pattern was unchanged. The autism main effect on gaze synchrony remained significant in all four contexts (β = −0.44 to −0.72, all q ≤ 0.005), and the ASD × ADHD interaction remained significant in the same three contexts as the primary models (TP: β = +0.43, q = 0.003; DM: β = +0.48, q = 0.001; WK: β = +0.40, q = 0.001), with *Fun with Fractals* again non-significant (β = +0.24, q = 0.176). The interaction is therefore not driven by a small number of extreme observations within the smallest cell. Absolute gaze phase was the exception: its autism main effects attenuated by roughly 60% under robust regression (DM: +0.41 to +0.17; WK: +0.35 to +0.12), and only the *Despicable Me* effect remained significant, indicating a sensitivity to extreme observations that the magnitude results do not share.

Finally, we examined whether severity or ascertainment differences between the autism-without-ADHD and comorbid groups could account for the interaction (Supplementary Table S17). The comorbid group showed *higher* autistic trait severity on the SRS Total (M = 70.13 vs 66.85, p = 0.011, d = 0.31), including on the restricted and repetitive behavior subscale (M = 70.64 vs 66.30, p = 0.003, d = 0.37), which is least susceptible to inflation by attentional symptoms. The groups did not differ in full-scale IQ (p = 0.390), age (p = 0.255) or anxiety diagnosis (p = 0.208). The comorbid group had higher CBCL Total scores (p = 0.006, d = 0.34) and a higher rate of oppositional/conduct diagnosis (18.0% vs 8.1%, p = 0.007), and carried on average 1.04 more diagnoses — a difference accounted for entirely by the ADHD diagnosis itself. Every clinical dimension on which the two groups differed was elevated in the comorbid group, which nonetheless showed the smaller reduction in synchrony.

## 4. Discussion

The present study used a large transdiagnostic pediatric cohort to map continuous and categorical dimensions of autism and ADHD onto gaze and pupillary synchrony during naturalistic viewing. Three findings emerged. First, autistic traits predicted reduced gaze synchrony across viewing contexts that differed widely in social and narrative content, whereas ADHD traits showed no association when modeled alone. Second, the clinical co-occurrence of ADHD was associated with a smaller, not larger, reduction in synchrony than autism alone, a non-additive pattern that survived adjustment for acquisition configuration, co-occurring psychopathology, stimulant exposure, and down-weighting of extreme observations. Third, coarse oculomotor measures computed from the identical recordings did not recover this structure. A fourth, methodological result is negative: the phase component of the complex correlation carried no information beyond its magnitude.

### 4.1. Reduction of synchrony with autistic traits across social and non-social contexts

Elevated autistic traits were associated with reduced gaze entrainment in the three videos with characters and narrative, and an autism diagnosis was associated with reduced entrainment in all four, including an abstract, non-social stimulus. This pattern is consistent with the social motivation account (Chevallier et al., 2012) but is not fully explained by it: a reduction confined to socially motivated attention would not be expected to appear during a geometric pattern sequence containing no faces, agents or narrative. The breadth of the effect suggests instead that autism is associated with a more general difference in how attention is allocated to a continuous dynamic visual stream, that is, a greater idiosyncrasy of attentional selection rather than reduced engagement with social content specifically. The two accounts are not exclusive. The reduction was smallest for the non-social stimulus, and its dimensional counterpart there fell short of significance, so social content may amplify a difference that is present without it. Pupillary synchrony showed the same direction with a narrower and less robust footprint: its dimensional associations were confined to two contexts and did not survive restriction to the categorical subsample.

### 4.2. Non-additive effects of co-occurring ADHD

Co-occurring autism and ADHD has frequently been characterized as a compounding of impairment, with greater difficulties than in either condition alone (Rao & Landa, 2014; Yerys et al., 2009). The present findings are not consistent with an additive model for this measure.

Autism in the absence of ADHD was associated with a large reduction in gaze synchrony relative to typically developing controls, whereas the corresponding reduction among children with co-occurring ADHD, relative to children with ADHD alone, was roughly two-thirds smaller. This sub-additive pattern appeared in three of the four viewing contexts in the full sample, and in all four when the analysis was restricted to a single acquisition configuration.

We characterize this pattern as an attenuated group difference rather than as a protective or compensatory effect, for two reasons. First, the designation of a co-occurring neurodevelopmental condition as protective is not warranted by a difference in a single physiological index, given that ADHD is independently associated with substantial functional impairment across academic, social and adaptive domains. Second, the present design is cross-sectional and observational, and permits no inference regarding the direction or mechanism of the association. The findings therefore bear on the physiological characterization of comorbid presentations rather than on their clinical prognosis.

The most obvious alternative explanation is that the comorbid group simply had milder autism. Three lines of evidence argue against it. First, parent-reported autistic trait severity was higher, not lower, in the comorbid group than in the group with autism alone. Second, this elevation extended to restricted and repetitive behavior, the SRS domain least susceptible to inflation by attentional and externalizing symptoms (Hus et al., 2013); an elevation confined to attention-loaded subscales would have pointed to instrument contamination rather than greater core severity. Third, every clinical dimension on which the two groups differed, including general psychopathology and oppositional or conduct diagnoses, was elevated in the comorbid group, which nevertheless showed the smaller reduction in synchrony. Confounding by clinical severity would therefore be expected to work against the observed interaction rather than to produce it, and consistent with this, the interaction grew stronger when those dimensions were statistically controlled.

One account of the mechanism is that the more frequent gaze shifting characteristic of ADHD interrupts the prolonged, idiosyncratic fixation associated with autism, incidentally increasing the overlap between an individual’s gaze path and the normative trajectory. Our data are consistent with the direction this account predicts but do not establish it. Comorbid children shifted gaze somewhat more often than children with autism alone in all four contexts, but the differences were small and none was statistically reliable. The mechanism should be regarded as a hypothesis the present data fail to refute, not one they confirm, and testing it properly requires oculomotor sampling finer than the 30 Hz common grid used here.

### 4.3. Dimensional and categorical frameworks diverge

The interaction appeared in the categorical models but not in the continuous trait models, where the product of autistic and ADHD traits predicted synchrony in no context. Two readings are available. The effect may be a genuine threshold phenomenon, emerging only when both conditions reach clinical severity, in which case categorical boundaries capture something dimensions do not. Alternatively, the dimensional null may reflect limited power: a product term formed from two noisy, substantially correlated parent-report instruments is poorly conditioned, and the absence of an interaction in such a model is weak evidence for its absence in the underlying phenotype. We cannot adjudicate between these with the present data, and note only that the frameworks are not interchangeable, which argues for reporting both rather than treating either as the definitive analysis.

The two frameworks were not wholly discordant, however. ADHD traits, which showed no association with gaze synchrony when modeled alone, predicted *higher* synchrony in one viewing context once autistic traits were held constant. This is the continuous analogues of the categorical interaction: in both frameworks, attentional symptoms accompany more normative gaze among children matched on autistic features. The result persisted, and strengthened, when the dimensional models were restricted to the participants used in the categorical analysis, so the divergence between the frameworks is not a consequence of their being fitted to different samples.

### 4.4. Absolute gaze phase does not index an independent phenomenon

We had expected the phase component of the complex correlation to provide information about the spatial geometry of misalignment that the modulus does not carry. It does not. Phase and magnitude shared a substantial portion of their variance, and the group differences observed in phase disappeared once the phase implied by each participant’s own magnitude was removed, whether that expectation was derived from simulation or estimated non-parametrically. The phase differences were also far more sensitive to extreme observations than the magnitude differences computed from the same recordings. The explanation is statistical rather than substantive: the argument of a complex correlation is estimated with precision that scales with its modulus, and gaze position is so strongly autocorrelated that the effective degrees of freedom available to the phase estimate are far fewer than the number of samples suggests.

We report this as a negative methodological result because the inference is not specific to these data. Any application of complex-correlation or phase-based synchrony to smooth, strongly autocorrelated signals is exposed to the same confound, and a magnitude-matched null is necessary before a phase difference between groups can be interpreted.

### 4.5. Relation to other gaze measures

Ten summary measures of oculomotor behavior derived from the same recordings, covering gaze-shift rate, dwell duration, dispersion, scan-path speed and central bias, did not recover the group structure that inter-subject synchrony detected. The single exception, longer dwell durations in autism without ADHD during one video, was not matched by its reciprocal measure, gaze-shift rate, and reflects a small number of long dwells rather than a general difference in viewing behavior. The scope of this null result should be stated carefully. These measures describe how the eyes move irrespective of what is on the screen: two children can shift gaze equally often and equally far while looking at different things. The result therefore indicates that the group differences lie in where gaze is directed in relation to the stimulus, not in the gross dynamics of eye movement. It does not show that inter-subject synchrony outperforms other content-referenced approaches, which we did not test.

Two such approaches are well established. The first quantifies fixation time on predefined regions of interest, typically faces, eyes and bodies, and underlies much of what is known about social attention in autism (Jones et al., 2023; Klin et al., 2002). The second scores gaze against a computational saliency model. Applying this approach to one of the videos used here, in the same cohort, Dziemian et al. (Dziemian et al., 2025) found that children with combined-presentation ADHD showed reduced correspondence with a saliency model trained on neurotypical viewing but not with a model of low-level visual features, and that basic fixation measures did not account for the difference, a pattern that parallels the dissociation reported here. Inter-subject synchrony belongs to the same family, with the typically developing group serving as an empirical reference in place of annotated regions or a model. This has practical advantages: it requires no annotation and no assumptions about which features matter, and it applies equally to a stimulus without faces or agents, for which social regions of interest are undefined. Its cost is interpretability, since a low synchrony value indicates that a child looked elsewhere without indicating where. Whether synchrony, region-of-interest and saliency-based measures capture the same variance, and whether the non-additive pattern reported here would appear in each, are open questions that the present recordings could address directly.

## 5. Limitations

### Precision and sample size

This was a secondary analysis of an existing cohort. The sample size was set by the number of usable recordings and not by a prior power calculation. Although the full sample is large, the group with autism and without ADHD is small, with 45 to 78 children per video, and this group limits the precision of every comparison that involves it. The interaction estimates are therefore imprecise: their 95% confidence intervals span roughly 0.6 to 0.7 standard deviations, so the size of the effect is known much less well than its direction.

### Replication

We did not test the findings in an independent cohort. The consistency of the gaze result across three videos and across analytic specifications is not a substitute for replication, because all analyses draw on the same children. The pupil interaction was observed in one video only and should be regarded as exploratory.

### Who the children were

Participants came from a help-seeking community cohort, and the findings may not generalize to other populations. Children who present to such a cohort with autism and no attentional diagnosis may be an unusual subgroup. Diagnostic overshadowing may make them atypical in ways we cannot assess, and this bears directly on the comparison at the center of the study.

### Diagnosis

Diagnoses were established by clinician consensus, without the standard research instruments for autism (ADOS-2, ADI-R). This constrains the categorical analyses in particular.

### Missing data

Children with higher autistic traits were less likely to provide usable recordings in three of the four videos. Because this narrows the range of traits in the analyzed sample, the reported trait associations are probably underestimates. Retention was markedly lower for *Fun with Fractals* than for the other videos, for reasons we could not establish; an abstract educational video may hold attention less reliably or may be less suitable for younger children.

### Acquisition

Eye-tracker configuration varied across recordings and was unevenly distributed across diagnostic groups. We adjusted for configuration and repeated the analysis within a single configuration, but the typically developing reference group could not itself be matched on configuration. The acquisition protocol is incompletely documented. According to the available documentation, *The Present* was always shown last, so differences between that video and the others cannot be separated from presentation order.

### Pupil measures

Pupil size responds to screen brightness. Synchrony in pupil size between children watching the same video therefore partly reflects shared changes in luminance, and the present design cannot separate these from shared cognitive or emotional responses.

### Medication

Medication status was documented for only about one fifth of participants, so an influence of medication cannot be excluded for the remainder.

### Design and clinical use

The design is cross-sectional and cannot show how these patterns develop or what causes them. Although the standardized group differences were large, the proportion of variance explained was small throughout. On the present evidence, these measures are not suitable for classifying individual children.

## 6. Conclusions

Autistic traits are associated with reduced gaze synchrony during naturalistic viewing, across social and non-social contexts alike. Co-occurring ADHD accompanies a smaller, not larger, reduction than autism alone, a pattern that withstood adjustment for acquisition, psychopathology and medication, was not driven by extreme observations, and was not recoverable from coarse oculomotor measures. A child with both conditions is therefore not simply a child with more of each: on this measure, the combination looks less like isolated autism, not more. These findings argue against additive models of autism–ADHD co-occurrence and for the use of model-free synchrony measures in transdiagnostic phenotyping, with the caution that the phase component of complex-correlation synchrony requires a magnitude-matched null before it can be interpreted.

## Abbreviations

ADHD: Attention-deficit/hyperactivity disorder
ADI-R: Autism Diagnostic Interview-Revised
ADOS-2: Autism Diagnostic Observation Schedule, Second Edition
ANCOVA: Analysis of covariance
ASD: Autism spectrum disorder
BCEA: Bivariate contour ellipse area
CBCL: Child Behavior Checklist
CI: Confidence interval
CR: Corneal reflection
DM: Despicable Me
FF: Fun with Fractals
HBN: Healthy Brain Network
ISC: Inter-subject correlation
K-SADS-COMP: Kiddie Schedule for Affective Disorders and Schizophrenia, Computerized version
MAD: Median absolute deviation
SD: Standard deviation
SRS-2: Social Responsiveness Scale, Second Edition
SWAN: Strengths and Weaknesses of ADHD Symptoms and Normal Behavior scale
TD: Typically developing
TP: The Present
WK: Diary of a Wimpy Kid

## Declarations

### Ethics approval and consent to participate

The data utilized in this study were obtained from the publicly available Healthy Brain Network (HBN) initiative. The original HBN study protocol was approved by the Chesapeake Institutional Review Board (now Advarra; Pro00012309), and all research was conducted in accordance with the Declaration of Helsinki. Prior to data collection, written informed consent was obtained from all adult participants and the legal guardians of all participants under 18 years of age. Written assent was additionally obtained from all participants under 18 years of age. The present study is a secondary analysis of de-identified data obtained under the Healthy Brain Network data usage agreement.

### Consent for publication

Not applicable.

### Availability of data and materials

The datasets analyzed during the current study are available through the Child Mind Institute’s Healthy Brain Network open-access initiative. The data can be accessed via the International Neuroimaging Data-sharing Initiative (INDI) at http://fcon_1000.projects.nitrc.org/indi/cmi_healthy_brain_network/. Eye-tracking recordings are openly accessible; phenotypic and diagnostic data are available to investigators who complete the Healthy Brain Network data usage agreement. The Python scripts used for eye-tracking preprocessing and statistical modeling are available from the corresponding author on reasonable request during the peer-review process, and will be made publicly available upon publication.

### Competing interests

The authors declare that they have no competing interests.

### Funding

This work was supported by the National Institutes of Health (NIH) under grant numbers R15MH125332 (to X.D.), 5R01MH131335 (to B.B.B.), and 1R01AG085665 (to B.B.B.). Additional support was provided by the New Jersey Governor’s Council for Medical Research and Treatment of Autism under grant number CAUT25BRP005 (to X.D.). The funding sources had no role in the study design, data collection, analysis, interpretation of the data, or decision to submit the manuscript for publication.

### Authors’ contributions

XD conceptualized the study, acquired funding, preprocessed and analyzed the eye-tracking data, conducted the statistical modeling, and drafted the manuscript. BBB acquired funding, provided methodological and physiological supervision, and critically reviewed and edited the manuscript. All authors read and approved the final manuscript.

## Acknowledgements

Not applicable.

---

**Additional file 1 (.docx).** Supplementary materials. Sections S1–S7, Figures S1–S5 and Tables S1–S20: sample composition and data retention, full dimensional model statistics, sensitivity analyses of the categorical models, absolute gaze phase, coarse oculomotor measures, severity comparison, and stimulant medication.

## Notes

### Competing Interest Statement

The authors have declared no competing interest.

### Summary of Updates

This version is a substantial revision. The title has been changed. Eye-tracking preprocessing was redone following published pupillometry guidelines, and differences in eye-tracker configuration across recordings are now documented and modeled in all analyses. The sample was updated (N = 2476). Inter-subject correlation is now computed against a typically developing reference of equal size for every participant. New analyses were added: a magnitude-matched null for gaze phase, ten coarse oculomotor measures, dimensional models on the categorical subsample, and sensitivity analyses for acquisition configuration, co-occurring psychopathology, stimulant medication and extreme observations. Confidence intervals are reported for the primary models. The main conclusion is unchanged: co-occurring ADHD is associated with a smaller reduction in gaze synchrony than autism alone. Gaze phase is no longer interpreted as an independent measure. The Introduction, Discussion and Limitations were rewritten, and the supplementary materials were reorganized.

