## Supplementary Materials for "Attenuated gaze synchrony in autism and its non-additive modification by co-occurring ADHD: a transdiagnostic eye-tracking study of naturalistic viewing"

**Contents**

**S1. Sample composition, data retention, and acquisition.** Flow of recordings through the pipeline; tests of whether data loss was independent of phenotype; the acquisition heterogeneity the main analyses carry as covariates; whether pupil-channel availability was phenotypically selected. *Figure S1; Tables S1–S2.*

**S2. Dimensional trait models.** Full statistics behind the dimensional models of main text Figure 2, the continuous SRS × SWAN interaction models, and the dimensional models refitted without the ‘Other’ group. *Figures S2–S3; Tables S3–S6.*

**S3. Sensitivity of the categorical models.** Four analyses testing whether the reported interaction depends on acquisition configuration, covariate choice, co-occurring psychopathology, or extreme observations. *Tables S7–S10.*

**S4. Absolute gaze phase.** Why the phase component of the complex correlation does not carry information beyond its magnitude, evaluated against both a simulated magnitude-matched null and a non-parametric residual. *Figure S4; Tables S11–S14.*

**S5. Coarse oculomotor measures.** Ten stimulus-independent gaze measures computed from the same recordings, and a directional test of the mechanism proposed for the interaction. *Figure S5; Tables S15–S16.*

**S6. Severity and ascertainment.** Whether the two autism groups differ on clinical dimensions that could account for the interaction. *Table S17.*

**S7. Stimulant medication.** Availability of same-day medication records, prevalence of active stimulant use, and models adjusted for it. *Tables S18–S20.*

**Conventions**

The following apply throughout unless a caption states otherwise, and are not repeated in individual captions.

Models are fitted separately within each of the four viewing paradigms, abbreviated and ordered throughout as *The Present* (TP), *Despicable Me* (DM), *Diary of a Wimpy Kid* (WK) and *Fun with Fractals* (FF). All models adjust for participant age, biological sex, tracker configuration (the four rate-by-correction combinations present in these data) and screen shape (4:3 against wider); pupil models additionally adjust for the proportion of missing pupil samples. Dependent variables and continuous predictors are standardized within paradigm, so coefficients are in standard-deviation units. Benjamini-Hochberg false discovery rate corrections are applied independently for each model term within each metric family across the four paradigms, and q denotes the corrected value. In the 2 × 2 models, autism and ADHD are entered as 0/1 indicators with their product, so the autism coefficient is the difference between the autism-without-ADHD group and typically developing controls, the ADHD coefficient is the difference between the ADHD-without-autism group and controls, and the interaction is the departure of the comorbid group from the sum of the two. Standardized group differences are Cohen’s *d*, the mean difference divided by the root mean square of the two group standard deviations.

Missing data are handled by listwise deletion applied per metric, so sample sizes differ across metrics within a table: a participant lacking a calibrated pupil channel contributes to the gaze analyses but not the pupil analyses. Dimensional models retain participants in the heterogeneous “Other” diagnostic category; categorical models exclude them. The two frameworks are therefore fitted to overlapping but different samples.

**Supplementary Section S1: Sample Composition, Data Retention, and Acquisition**

Eye-tracking data in the Healthy Brain Network were acquired over an extended multi-site collection in which hardware configuration was not held constant, and recordings were lost at several stages of preprocessing. This section documents the flow of recordings through the pipeline, tests whether the losses were independent of phenotype, and characterizes the acquisition heterogeneity that the main analyses carry as covariates.

**S1.1. Participant flow**

Supplementary Figure S1 gives the flow of recordings through acquisition, preprocessing and clinical characterization, with exclusion counts at each stage for each of the four viewing paradigms. Of the recordings identified, 77.6% (*The Present*), 72.4% (*Despicable Me*), 75.3% (*Diary of a Wimpy Kid*) and 54.5% (*Fun with Fractals*) survived preprocessing. The lower yield for *Fun with Fractals* reflects exclusion rather than availability: a comparable number of recordings was identified for that paradigm (n = 2617) as for *The Present* (n = 2729). Requiring complete clinical characterization and covariates reduced these to the analytic samples of 1775, 1959, 2042 and 1186 reported in Table 1.

**Supplementary Figure S1.** Flow of eye-tracking recordings through preprocessing and analysis. Counts are recordings; a participant contributes up to four, one per viewing paradigm. Criteria are applied in the order shown. The four terminal samples differ on two independent restrictions: channel, since a calibrated pupil diameter exists only for corneal-reflection corrected recordings (1083, 1176, 1308 and 545 of the gaze analytic sample respectively), and diagnostic scope, since the dimensional models retain the heterogeneous “Other” group while the categorical models exclude it. The dimensional samples are further reduced by the requirement for complete SRS-2 and SWAN reports. All four are nested within the gaze analytic sample, and are counted with the same listwise deletion the corresponding models use, so they match the sample sizes of main text Table 2 and Supplementary Tables S3 and S4.

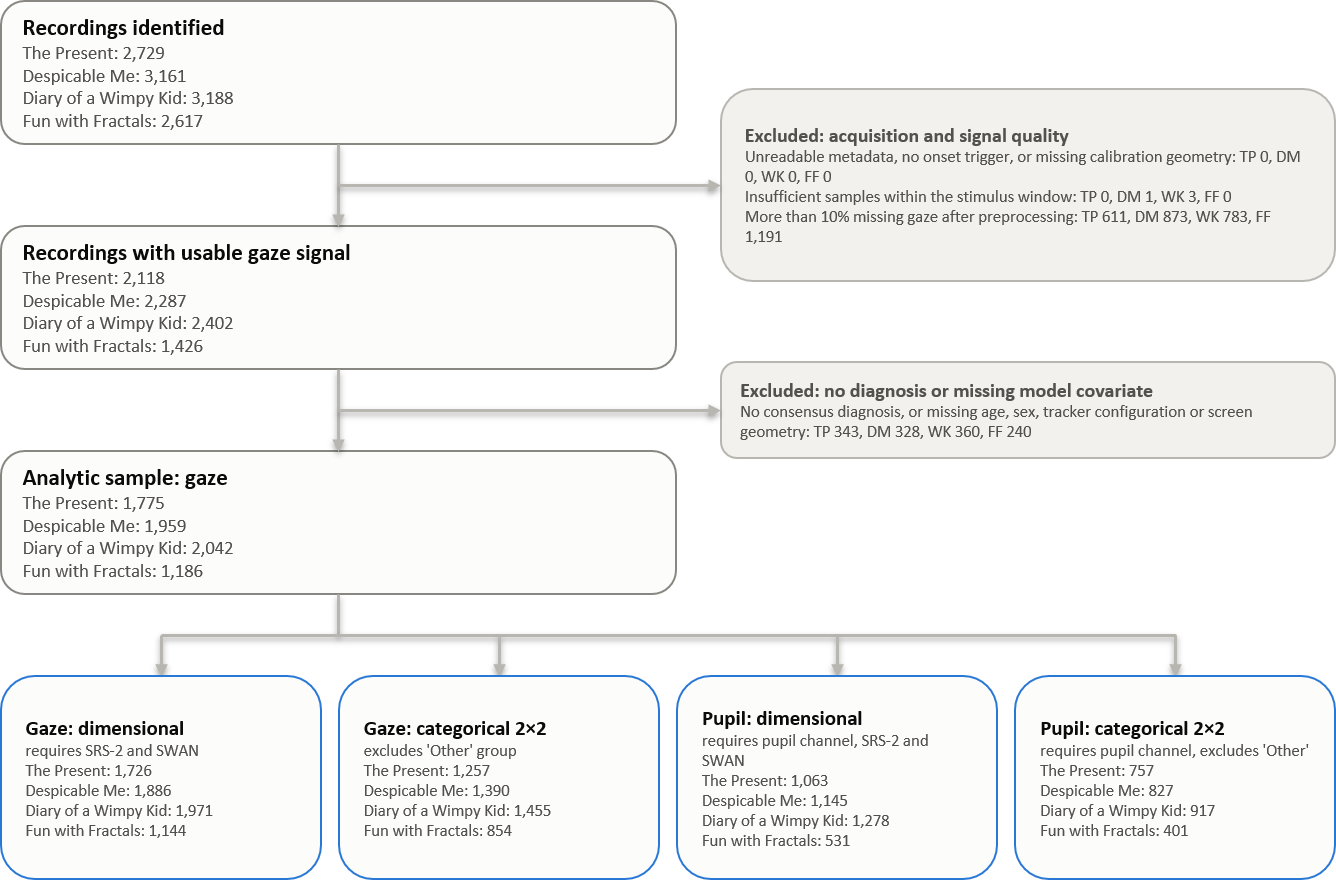

**S1.2. Retention was not independent of phenotype**

Because recordings were not lost at random, we modeled retention directly. Per-video logistic regressions of retention on SRS, SWAN, age and sex (Supplementary Table S1) show that higher autistic traits predicted lower retention in three of the four paradigms (*Despicable Me*: b = −0.011, p = 0.013; *Diary of a Wimpy Kid*: b = −0.016, p = 0.0003; *Fun with Fractals*: b = −0.015, p = 0.0007), with no association for *The Present* (p = 0.16). ADHD traits predicted retention in no paradigm (all p > 0.11).

**Supplementary Table S1.** Per-video logistic regression of recording retention on parent-reported autistic traits (SRS-2 Total T), ADHD traits (SWAN Total), age and sex, over all identified recordings with complete phenotype. Percentages retained are computed over recordings with complete phenotype and therefore differ slightly from the preprocessing yields in Section S1.1, which are computed over all identified recordings.

| Video | n | % retained | Term | coef | se | z | p |
| --- | --- | --- | --- | --- | --- | --- | --- |
| TP | 2261 | 76.4 | Autistic traits (SRS-2 Total T) | -0.007 | 0.005 | -1.397 | 0.162 |
|  |  |  | ADHD traits (SWAN Total) | -0.056 | 0.061 | -0.920 | 0.358 |
|  |  |  | Age (years) | -0.033 | 0.015 | -2.181 | 0.029 |
|  |  |  | Sex (female vs male) | 0.065 | 0.107 | 0.613 | 0.540 |
| DM | 2637 | 71.5 | Autistic traits (SRS-2 Total T) | -0.011 | 0.004 | -2.487 | 0.013 |
|  |  |  | ADHD traits (SWAN Total) | -0.017 | 0.051 | -0.338 | 0.735 |
|  |  |  | Age (years) | -0.028 | 0.013 | -2.108 | 0.035 |
|  |  |  | Sex (female vs male) | 0.073 | 0.093 | 0.786 | 0.432 |
| WK | 2647 | 74.5 | Autistic traits (SRS-2 Total T) | -0.016 | 0.004 | -3.577 | < 0.001 |
|  |  |  | ADHD traits (SWAN Total) | 0.084 | 0.053 | 1.584 | 0.113 |
|  |  |  | Age (years) | -0.005 | 0.014 | -0.395 | 0.693 |
|  |  |  | Sex (female vs male) | 0.127 | 0.096 | 1.322 | 0.186 |
| FF | 2121 | 53.9 | Autistic traits (SRS-2 Total T) | -0.015 | 0.004 | -3.385 | 0.001 |
|  |  |  | ADHD traits (SWAN Total) | -0.013 | 0.051 | -0.244 | 0.807 |
|  |  |  | Age (years) | 0.019 | 0.014 | 1.397 | 0.163 |
|  |  |  | Sex (female vs male) | 0.060 | 0.093 | 0.640 | 0.522 |

This has a direction. Children with higher autistic traits are under-represented among retained recordings, which restricts the range of the predictor in exactly the analyses that estimate trait associations. The reported dimensional effects are therefore likely to be conservative rather than inflated, and the per-video samples are not interchangeable.

**S1.3. Acquisition configuration was unbalanced across diagnostic groups**

Tracker configuration, sampling rate and display geometry all varied across recordings (main text Section 2.3). Chi-square tests of each acquisition variable against diagnostic group, within each video, are given in Supplementary Table S2. Tracker configuration, sampling rate, display resolution and screen aspect ratio were unbalanced across diagnostic groups in all four paradigms (all p ≤ 0.035; tracker configuration p ≤ 4.9 × 10⁻⁵ throughout). The associations were statistically reliable but small: Cramér’s V ranged from 0.084 to 0.129 across these tests, so acquisition explains a minor share of the variation in group membership while being reliably non-random with respect to it. This is why all models carry tracker configuration and screen shape as covariates, and why the configuration-restricted analysis of Table S7 is reported.

**Supplementary Table S2.** Chi-square tests of acquisition variables against diagnostic group, within each viewing paradigm. Aspect Ratio is the width-to-height ratio of the calibrated display area (three values). Sample Rate is the nominal sampling rate (30, 60 or 120 Hz). Has CR indicates whether corneal-reflection correction was active, and Pupil Usable whether the recording yielded a pupil channel that passed quality control. Tracker is the four-level combination of sampling rate and corneal-reflection correction entered in the models, and Wide Screen distinguishes 4:3 from wider calibrated areas. Config is Tracker crossed with Wide Screen (six combinations present). Cramér’s V is reported alongside p, since with samples of 1186–2042 a small association reaches significance.

| Variable | Video | Χ^2^ | df | p | Cramér’s V |
| --- | --- | --- | --- | --- | --- |
| Aspect Ratio | TP | 30.826 | 8 | < 0.001 | 0.093 |
|  | DM | 32.982 | 8 | < 0.001 | 0.092 |
|  | WK | 30.279 | 8 | < 0.001 | 0.086 |
|  | FF | 16.562 | 8 | 0.035 | 0.084 |
| Config | TP | 72.970 | 20 | < 0.001 | 0.101 |
|  | DM | 90.032 | 20 | < 0.001 | 0.107 |
|  | WK | 78.036 | 20 | < 0.001 | 0.098 |
|  | FF | 50.781 | 20 | < 0.001 | 0.103 |
| Has CR | TP | 14.750 | 4 | 0.005 | 0.091 |
|  | DM | 5.939 | 4 | 0.204 | 0.055 |
|  | WK | 4.939 | 4 | 0.294 | 0.049 |
|  | FF | 5.002 | 4 | 0.287 | 0.065 |
| Pupil Usable | TP | 12.674 | 4 | 0.013 | 0.084 |
|  | DM | 6.193 | 4 | 0.185 | 0.056 |
|  | WK | 5.542 | 4 | 0.236 | 0.052 |
|  | FF | 3.171 | 4 | 0.530 | 0.052 |
| Sample Rate | TP | 30.253 | 8 | < 0.001 | 0.092 |
|  | DM | 33.909 | 8 | < 0.001 | 0.093 |
|  | WK | 36.213 | 8 | < 0.001 | 0.094 |
|  | FF | 23.009 | 8 | 0.003 | 0.098 |
| Tracker | TP | 59.341 | 12 | < 0.001 | 0.106 |
|  | DM | 69.804 | 12 | < 0.001 | 0.109 |
|  | WK | 63.623 | 12 | < 0.001 | 0.102 |
|  | FF | 41.011 | 12 | < 0.001 | 0.107 |
| Wide Screen | TP | 29.685 | 4 | < 0.001 | 0.129 |
|  | DM | 31.383 | 4 | < 0.001 | 0.127 |
|  | WK | 27.878 | 4 | < 0.001 | 0.117 |
|  | FF | 10.858 | 4 | 0.028 | 0.096 |

Corneal-reflection correction is the exception. Its availability did not differ across diagnostic groups in *Despicable Me* (p = 0.204), *Diary of a Wimpy Kid* (p = 0.294) or *Fun with Fractals* (p = 0.287), and differed only in *The Present* (p = 0.0052, V = 0.091). The same pattern held for the downstream availability of a usable pupil channel (*The Present* p = 0.013; all others p ≥ 0.185).

**S1.4. Pupil-channel availability was not phenotypically selected**

Approximately 30% of recordings were made on a configuration without corneal-reflection correction and therefore carry no calibrated pupil diameter, which accounts for roughly four fifths of the difference between the gaze and pupil sample sizes in Table 1. Because this determines which participants contribute to the pupil analyses, we tested whether it was related to phenotype.

Among recordings that passed the gaze pipeline, we regressed corneal-reflection availability, and separately the availability of a usable pupil channel, on autistic traits, ADHD traits, age and sex within each paradigm. Across the sixteen trait tests one reached nominal significance — autistic traits in *The Present*, OR 0.877 per SD, p = 0.023 — which does not survive correction across those tests (smallest q = 0.36), and signal quality within corrected recordings was unrelated to either trait domain (all p ≥ 0.24). Pupil-channel availability was, however, associated with age in three of four paradigms (b = −0.064 to −0.043, p ≤ 0.004), older children being less likely to have been recorded on a corrected configuration; this is most plausibly a period effect, since collection spanned years over which configurations and referral composition both changed. Age is a covariate in all models. The nominal result in *The Present* converges with the acquisition imbalance reported for the same paradigm in S1.3.

**Supplementary Section S2: Dimensional Trait Models**

The main text reports the standardized coefficients of the dimensional models in Figure 2 (Section 3.3). This section gives the full model statistics behind them, the continuous interaction models, and the models refitted without the ‘Other’ group.

**S2.1. Full statistics for the single-trait and joint models**

Supplementary Tables S3 and S4 report, for every metric and viewing context, the standardized coefficient, partial eta-squared, uncorrected p, Benjamini-Hochberg q, and sample size for the models summarized in panels A and B of main text Figure 2 respectively. All models adjust for age, sex, tracker configuration and screen shape, with dependent variables and trait scores standardized within video.

**Supplementary Table S3.** Full statistics for the single-trait dimensional models summarized in panel A of main text Figure 2. Each trait domain is entered in its own model, adjusted for age, sex, tracker configuration and screen shape. Benjamini-Hochberg q values are computed within each metric family across the four viewing contexts.

| Metric | Term | Video | n | β | η_p_^2^ | p | q |
| --- | --- | --- | --- | --- | --- | --- | --- |
| Gaze  ISC  Magnitude | SRS-2 Total T | TP | 1726 | -0.116 | 0.014 | < 0.001 | < 0.001 |
|  | SRS-2 Total T | DM | 1886 | -0.069 | 0.005 | 0.003 | 0.004 |
|  | SRS-2 Total T | WK | 1971 | -0.095 | 0.009 | < 0.001 | < 0.001 |
|  | SRS-2 Total T | FF | 1144 | -0.055 | 0.003 | 0.061 | 0.061 |
|  | SWAN Total | TP | 1726 | 0.015 | 0.000 | 0.557 | 0.743 |
|  | SWAN Total | DM | 1886 | 0.004 | 0.000 | 0.859 | 0.859 |
|  | SWAN Total | WK | 1971 | -0.048 | 0.002 | 0.036 | 0.142 |
|  | SWAN Total | FF | 1144 | -0.041 | 0.002 | 0.184 | 0.368 |
| Absolute  Gaze  Phase | SRS-2 Total T | TP | 1726 | 0.005 | 0.000 | 0.833 | 0.877 |
|  | SRS-2 Total T | DM | 1886 | -0.004 | 0.000 | 0.877 | 0.877 |
|  | SRS-2 Total T | WK | 1971 | 0.014 | 0.000 | 0.530 | 0.877 |
|  | SRS-2 Total T | FF | 1144 | -0.031 | 0.001 | 0.291 | 0.877 |
|  | SWAN Total | TP | 1726 | 0.003 | 0.000 | 0.902 | 0.956 |
|  | SWAN Total | DM | 1886 | -0.021 | 0.000 | 0.384 | 0.956 |
|  | SWAN Total | WK | 1971 | 0.001 | 0.000 | 0.956 | 0.956 |
|  | SWAN Total | FF | 1144 | -0.006 | 0.000 | 0.860 | 0.956 |
| Pupil  ISC | SRS-2 Total T | TP | 1063 | -0.087 | 0.008 | 0.004 | 0.017 |
|  | SRS-2 Total T | DM | 1145 | -0.047 | 0.002 | 0.112 | 0.149 |
|  | SRS-2 Total T | WK | 1278 | -0.072 | 0.005 | 0.011 | 0.022 |
|  | SRS-2 Total T | FF | 531 | -0.055 | 0.003 | 0.206 | 0.206 |
|  | SWAN Total | TP | 1063 | 0.022 | 0.001 | 0.475 | 0.950 |
|  | SWAN Total | DM | 1145 | -0.002 | 0.000 | 0.960 | 0.960 |
|  | SWAN Total | WK | 1278 | -0.039 | 0.001 | 0.184 | 0.735 |
|  | SWAN Total | FF | 531 | -0.009 | 0.000 | 0.848 | 0.960 |

**Supplementary Table S4.** Full statistics for the two-trait dimensional models summarized in panel B of main text Figure 2, in which both trait domains are entered simultaneously so that each coefficient reflects the association net of the other.

| Metric | Term | Video | n | β | η_p_^2^ | p | q |
| --- | --- | --- | --- | --- | --- | --- | --- |
| Gaze  ISC  Magnitude | SRS-2 Total T | TP | 1726 | -0.158 | 0.020 | < 0.001 | < 0.001 |
|  | SRS-2 Total T | DM | 1886 | -0.092 | 0.007 | < 0.001 | 0.001 |
|  | SRS-2 Total T | WK | 1971 | -0.093 | 0.007 | < 0.001 | < 0.001 |
|  | SRS-2 Total T | FF | 1144 | -0.047 | 0.002 | 0.159 | 0.159 |
|  | SWAN Total | TP | 1726 | 0.092 | 0.006 | 0.001 | 0.004 |
|  | SWAN Total | DM | 1886 | 0.051 | 0.002 | 0.063 | 0.126 |
|  | SWAN Total | WK | 1971 | -0.003 | 0.000 | 0.905 | 0.905 |
|  | SWAN Total | FF | 1144 | -0.017 | 0.000 | 0.629 | 0.838 |
| Absolute  Gaze  Phase | SRS-2 Total T | TP | 1726 | 0.005 | 0.000 | 0.863 | 0.863 |
|  | SRS-2 Total T | DM | 1886 | 0.008 | 0.000 | 0.761 | 0.863 |
|  | SRS-2 Total T | WK | 1971 | 0.017 | 0.000 | 0.495 | 0.863 |
|  | SRS-2 Total T | FF | 1144 | -0.038 | 0.001 | 0.268 | 0.863 |
|  | SWAN Total | TP | 1726 | 0.001 | 0.000 | 0.979 | 0.979 |
|  | SWAN Total | DM | 1886 | -0.025 | 0.000 | 0.363 | 0.979 |
|  | SWAN Total | WK | 1971 | -0.007 | 0.000 | 0.786 | 0.979 |
|  | SWAN Total | FF | 1144 | 0.014 | 0.000 | 0.703 | 0.979 |
| Pupil  ISC | SRS-2 Total T | TP | 1063 | -0.120 | 0.012 | < 0.001 | 0.002 |
|  | SRS-2 Total T | DM | 1145 | -0.059 | 0.003 | 0.078 | 0.103 |
|  | SRS-2 Total T | WK | 1278 | -0.069 | 0.004 | 0.029 | 0.058 |
|  | SRS-2 Total T | FF | 531 | -0.066 | 0.003 | 0.184 | 0.184 |
|  | SWAN Total | TP | 1063 | 0.076 | 0.005 | 0.027 | 0.109 |
|  | SWAN Total | DM | 1145 | 0.027 | 0.001 | 0.439 | 0.867 |
|  | SWAN Total | WK | 1278 | -0.005 | 0.000 | 0.874 | 0.874 |
|  | SWAN Total | FF | 531 | 0.024 | 0.000 | 0.650 | 0.867 |

**Supplementary Figure S2.** Autistic traits (SRS-2 Total T) against each synchrony metric, by video. Rows: gaze ISC magnitude, absolute gaze phase and pupil ISC. Columns: the four videos. Each point is one participant; the line is the ordinary least-squares fit with its 95% confidence band, and the Pearson correlation and *n* are printed in each panel. Vertical axes are clipped to the central 99% of each metric for legibility; fits use all data. The full cohort is included, the ‘Other’ diagnostic group among them. Estimates adjusted for age, sex, tracker configuration and screen shape are given in Tables S3 and S4; the fits drawn here are unadjusted.

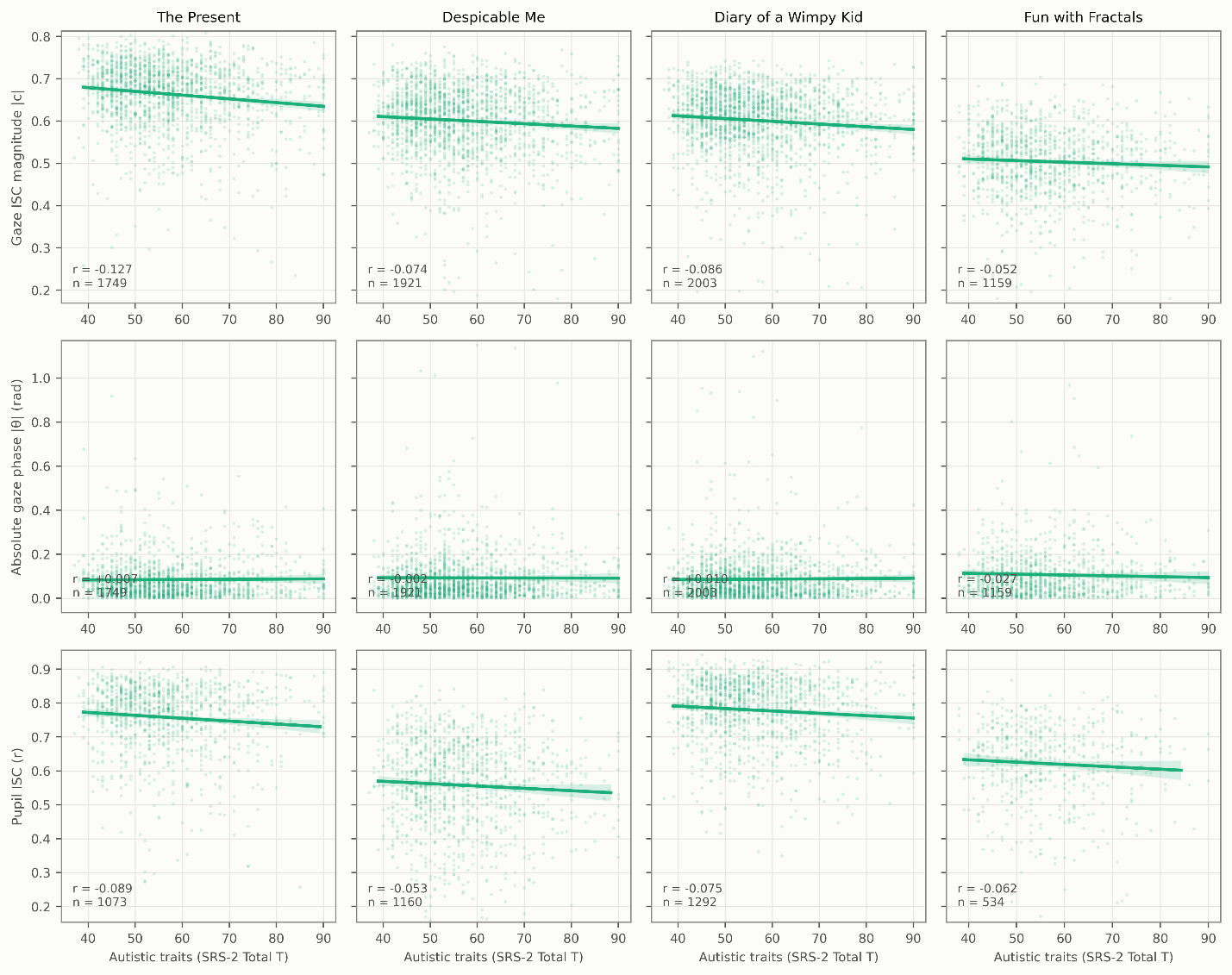

**Supplementary Figure S3.** ADHD traits (SWAN Total) against each synchrony metric, by video. Rows: gaze ISC magnitude, absolute gaze phase and pupil ISC. Columns: the four videos. Each point is one participant; the line is the ordinary least-squares fit with its 95% confidence band, and the Pearson correlation and *n* are printed in each panel. Vertical axes are clipped to the central 99% of each metric for legibility; fits use all data. The full cohort is included, the ‘Other’ diagnostic group among them. Estimates adjusted for age, sex, tracker configuration and screen shape are given in Tables S3 and S4; the fits drawn here are unadjusted.

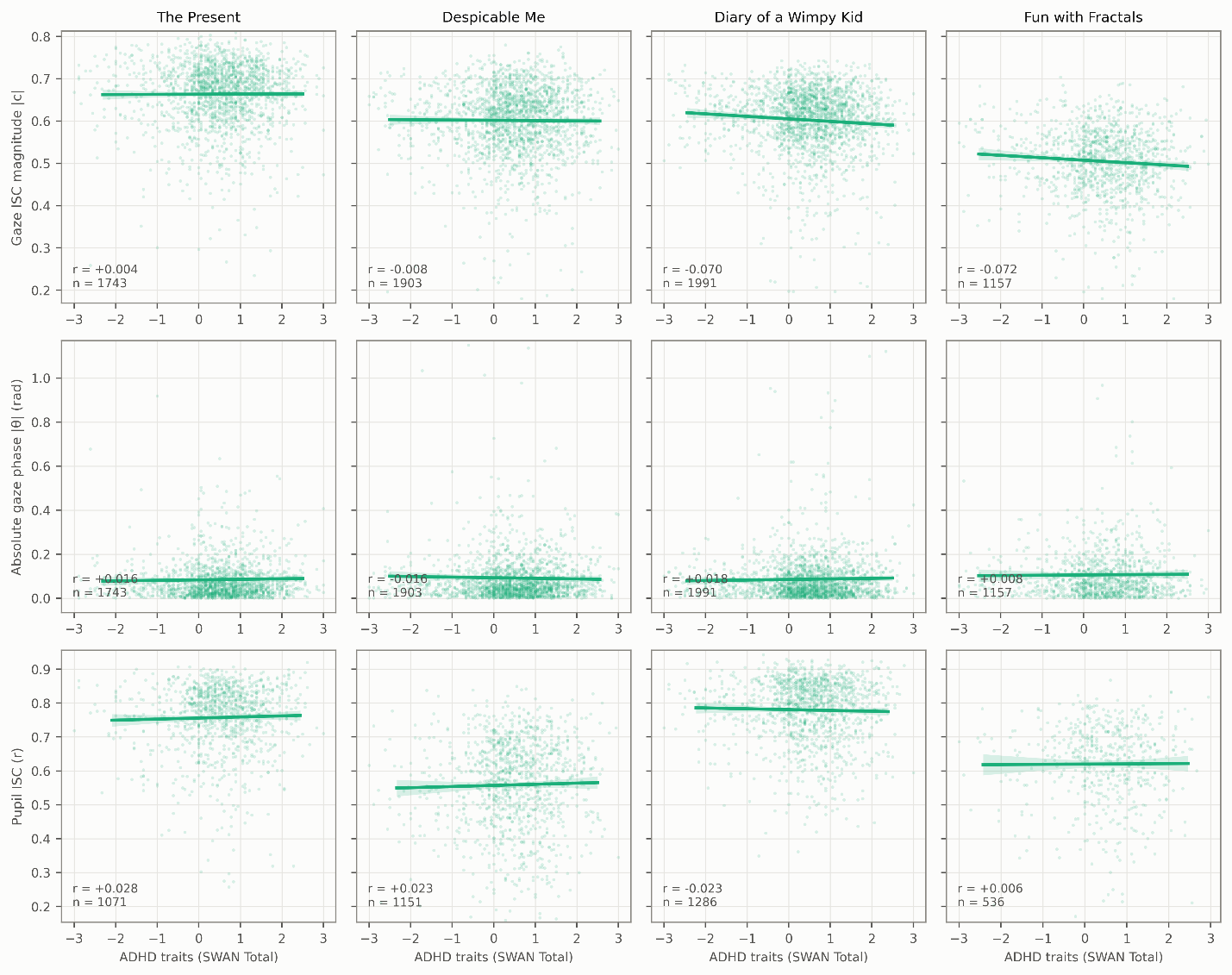

These models retain participants in the heterogeneous “Other” diagnostic category and require complete SRS and SWAN reports, giving samples of 1144–1971 for gaze metrics and 531–1278 for pupil ISC. The categorical models of main text Table 2 exclude the “Other” group and so run on 854–1455 participants. The two frameworks are therefore fitted to overlapping but different samples, which is one reason their results are not directly interchangeable.

**S2.2. Continuous trait interactions**

To test whether highly comorbid presentations compound dimensionally, as the categorical interaction suggests they do not, we added an SRS × SWAN product term to the joint models. Results are given in Supplementary Table S5.

**Supplementary Table S5.** Continuous SRS × SWAN interaction terms added to the joint dimensional models of Table S4. Standardized coefficients, partial eta-squared, uncorrected p and Benjamini-Hochberg q within each metric family across the four viewing contexts. Models adjust for age, sex, tracker configuration and screen shape, and include participants in the “Other” diagnostic category.

| Metric | Video | n | β | η_p_^2^ | p | q |
| --- | --- | --- | --- | --- | --- | --- |
| Gaze  ISC  Magnitude | TP | 1726 | 0.016 | 0.000 | 0.460 | 0.460 |
|  | DM | 1886 | 0.035 | 0.001 | 0.105 | 0.211 |
|  | WK | 1971 | 0.028 | 0.001 | 0.172 | 0.230 |
|  | FF | 1144 | 0.045 | 0.002 | 0.104 | 0.211 |
| Absolute  Gaze  Phase | TP | 1726 | 0.024 | 0.001 | 0.294 | 0.392 |
|  | DM | 1886 | -0.017 | 0.000 | 0.433 | 0.433 |
|  | WK | 1971 | -0.036 | 0.002 | 0.087 | 0.327 |
|  | FF | 1144 | -0.039 | 0.002 | 0.163 | 0.327 |
| Pupil  ISC | TP | 1063 | 0.033 | 0.001 | 0.244 | 0.964 |
|  | DM | 1145 | 0.001 | 0.000 | 0.964 | 0.964 |
|  | WK | 1278 | 0.003 | 0.000 | 0.926 | 0.964 |
|  | FF | 531 | -0.020 | 0.000 | 0.651 | 0.964 |

No interaction term approached significance in any metric or context. Coefficients were uniformly small (|β| ≤ 0.045) and partial eta-squared values negligible (≤ 0.0023); the smallest corrected q was 0.21. For gaze ISC magnitude the four coefficients were +0.016 (*The Present*), +0.035 (*Despicable Me*), +0.028 (*Diary of a Wimpy Kid*) and +0.045 (*Fun with Fractals*).

Two features of this null deserve comment, because the contrast with the significant categorical interaction invites a stronger reading than the data support.

First, all four gaze coefficients carry the same sign as the categorical interaction. A pure threshold account — in which the attenuation emerges only once both conditions reach clinical severity — predicts no systematic signal in the sub-clinical range, not a consistently positive one. The four contexts draw on largely the same participants and so are correlated rather than independent, and none of the estimates is close to significance, so this is suggestive only; but it is suggestive in the direction of limited power rather than of a genuine discontinuity.

Second, these product terms are poorly conditioned. SRS and SWAN totals correlate r = 0.46–0.47 in this sample, so their product is substantially collinear with both constituent main effects, and both instruments are noisy parent reports whose measurement error compounds multiplicatively in the product. The absence of an interaction in models of this form is weak evidence for the absence of an interaction in the underlying phenotype.

We therefore report the dimensional and categorical frameworks as giving different answers without adjudicating between them (main text Section 4.3), rather than treating the dimensional null as establishing that the attenuation requires categorical comorbidity.

**S2.3. Dimensional models excluding the “Other” group**

The dimensional models retain the heterogeneous “Other” diagnostic category while the categorical models exclude it, so the two frameworks are fitted to overlapping but different samples. To establish that their divergence is not an artifact of that difference, we repeated the dimensional models on the categorical subsample (Table S6).

The gaze findings are essentially unchanged. The single-trait autistic coefficient was −0.116, −0.081, −0.093 and −0.057 against −0.116, −0.069, −0.095 and −0.055 in the full cohort — unchanged to three decimal places in *The Present*, marginally stronger in *Despicable Me*, and otherwise indistinguishable. Significance was retained in the same three contexts despite the loss of roughly 29% of participants, with *Fun with Fractals* again falling short (q = 0.100 against 0.061). The joint models likewise strengthened slightly rather than attenuating, the autistic-trait coefficient moving from −0.158 to −0.168 in *The Present* and from −0.092 to −0.103 in *Despicable Me*.

The suppression effect is more pronounced on this sample, not less. In *The Present*, the ADHD-trait coefficient rose from +0.092 (q = 0.004) to +0.117 (q = 0.001) once autistic traits were held constant — so the continuous counterpart of the categorical interaction is present, and strengthened, in precisely the participants the categorical models use. Whatever separates the two frameworks, it is not sample composition.

Absolute gaze phase remained null throughout, under both specifications and in every context (all q ≥ 0.80).

The pupil results do not survive the restriction. The single-trait autistic coefficient attenuated from −0.087 to −0.069 in *The Present* and from −0.072 to −0.054 in *Diary of a Wimpy Kid*, and neither remained significant (both q = 0.182 against 0.017 and 0.022). The pupil samples lose about 30% of their participants under this restriction — 1063 to 742 and 1278 to 896 — so the loss of significance is partly a loss of power, but the coefficients themselves shrink by roughly a fifth, which power alone does not explain. We therefore report the pupil dimensional associations as holding in the full referred sample but not demonstrable within the categorical subsample.

**Supplementary Table S6.** Dimensional models refitted with the heterogeneous ‘Other’ diagnostic group excluded, so the dimensional and categorical frameworks are estimated on the same participants. Each trait entered alone (sep) on the left, both entered together (joint) on the right.

| Metric | Video | n | β_SRS _sep_ | q_SRS_sep_ | β_SWAN _sep_ | q_SWAN_sep_ | β_SRS_ joint_ | q_SRS_ joint_ | β_SWAN_ joint_ | q_SWAN_ joint_ |
| --- | --- | --- | --- | --- | --- | --- | --- | --- | --- | --- |
| Gaze  ISC  Magnitude | TP | 1223 | -0.116 | < 0.001 | 0.039 | 0.537 | -0.168 | < 0.001 | 0.117 | 0.001 |
|  | DM | 1341 | -0.081 | 0.004 | -0.001 | 0.974 | -0.103 | 0.001 | 0.049 | 0.256 |
|  | WK | 1408 | -0.093 | 0.001 | -0.030 | 0.537 | -0.100 | 0.001 | 0.016 | 0.790 |
|  | FF | 825 | -0.057 | 0.100 | -0.029 | 0.551 | -0.055 | 0.153 | -0.004 | 0.922 |
| Absolute  Gaze  Phase | TP | 1223 | 0.010 | 0.805 | 0.001 | 0.960 | 0.011 | 0.927 | -0.004 | 0.909 |
|  | DM | 1341 | -0.007 | 0.805 | -0.020 | 0.960 | 0.003 | 0.927 | -0.021 | 0.909 |
|  | WK | 1408 | 0.021 | 0.805 | 0.006 | 0.960 | 0.022 | 0.908 | -0.004 | 0.909 |
|  | FF | 825 | -0.037 | 0.805 | -0.008 | 0.960 | -0.042 | 0.908 | 0.011 | 0.909 |
| Pupil  ISC | TP | 742 | -0.069 | 0.182 | 0.025 | 0.819 | -0.096 | 0.064 | 0.066 | 0.420 |
|  | DM | 806 | -0.047 | 0.182 | -0.008 | 0.819 | -0.054 | 0.168 | 0.017 | 0.908 |
|  | WK | 896 | -0.054 | 0.182 | -0.028 | 0.819 | -0.052 | 0.168 | -0.004 | 0.908 |
|  | FF | 392 | -0.075 | 0.182 | -0.023 | 0.819 | -0.081 | 0.168 | 0.013 | 0.908 |

**Supplementary Section S3: Sensitivity of the Categorical Models**

The interaction reported in the main text (main text Table 2 and Figure 3) is estimated from a design in which the smallest cell contains 45–78 participants per context and in which acquisition configuration is unbalanced across diagnostic groups (Section S1.3). This section reports four analyses testing whether the result depends on those features. Main text Figure 5 displays the interaction estimate under each covariate and sample specification, including the medication-adjusted model of Section S7.

**S3.1. Restriction to a single acquisition configuration**

Tracker configuration was unbalanced across diagnostic groups in every paradigm, and the typically developing reference could not be configuration-matched, since some strata contain fewer than ten typically developing participants per video. Covariate adjustment is therefore not a complete remedy. We repeated the models within the single dominant configuration, 60 Hz with corneal-reflection correction (Supplementary Table S7).

**Supplementary Table S7.** The 2 × 2 analysis of covariance restricted to recordings made at 60 Hz with corneal-reflection correction, the single dominant acquisition configuration. Tracker configuration is unbalanced across diagnostic groups (Table S2) and the typically developing reference cannot be configuration-matched, so this restriction is the configuration-homogeneous check that covariate adjustment cannot provide. Cell sizes per paradigm (TD / ASD only / ADHD only / comorbid): TP 76/33/454/90; DM 108/40/492/107; WK 116/48/543/113; FF 77/20/284/57.

| Metric | Term | Video | n | β | η_p_^2^ | p | q |
| --- | --- | --- | --- | --- | --- | --- | --- |
| Gaze  ISC  Magnitude | ASD | TP | 653 | -1.153 | 0.047 | < 0.001 | < 0.001 |
|  | ASD | DM | 747 | -1.007 | 0.039 | < 0.001 | < 0.001 |
|  | ASD | WK | 820 | -0.853 | 0.031 | < 0.001 | < 0.001 |
|  | ASD | FF | 438 | -0.864 | 0.027 | 0.001 | 0.001 |
|  | ADHD | TP | 653 | -0.220 | 0.005 | 0.071 | 0.141 |
|  | ADHD | DM | 747 | -0.146 | 0.003 | 0.168 | 0.223 |
|  | ADHD | WK | 820 | -0.093 | 0.001 | 0.352 | 0.352 |
|  | ADHD | FF | 438 | -0.229 | 0.008 | 0.071 | 0.141 |
|  | ASD × ADHD | TP | 653 | 1.008 | 0.028 | < 0.001 | < 0.001 |
|  | ASD × ADHD | DM | 747 | 0.838 | 0.021 | < 0.001 | < 0.001 |
|  | ASD × ADHD | WK | 820 | 0.679 | 0.015 | 0.001 | 0.001 |
|  | ASD × ADHD | FF | 438 | 0.679 | 0.013 | 0.018 | 0.018 |
| Absolute  Gaze  Phase | ASD | TP | 653 | 0.127 | 0.001 | 0.543 | 0.543 |
|  | ASD | DM | 747 | 0.343 | 0.005 | 0.065 | 0.130 |
|  | ASD | WK | 820 | 0.682 | 0.019 | < 0.001 | < 0.001 |
|  | ASD | FF | 438 | 0.319 | 0.004 | 0.213 | 0.284 |
|  | ADHD | TP | 653 | 0.043 | 0.000 | 0.728 | 0.754 |
|  | ADHD | DM | 747 | 0.060 | 0.000 | 0.573 | 0.754 |
|  | ADHD | WK | 820 | 0.145 | 0.003 | 0.158 | 0.632 |
|  | ADHD | FF | 438 | 0.041 | 0.000 | 0.754 | 0.754 |
|  | ASD × ADHD | TP | 653 | 0.066 | 0.000 | 0.781 | 0.781 |
|  | ASD × ADHD | DM | 747 | -0.245 | 0.002 | 0.250 | 0.334 |
|  | ASD × ADHD | WK | 820 | -0.494 | 0.008 | 0.013 | 0.054 |
|  | ASD × ADHD | FF | 438 | -0.370 | 0.004 | 0.208 | 0.334 |
| Pupil  ISC | ASD | TP | 606 | -0.932 | 0.032 | < 0.001 | < 0.001 |
|  | ASD | DM | 671 | -0.478 | 0.009 | 0.013 | 0.026 |
|  | ASD | WK | 750 | -0.389 | 0.006 | 0.029 | 0.039 |
|  | ASD | FF | 365 | -0.127 | 0.001 | 0.652 | 0.652 |
|  | ADHD | TP | 606 | -0.083 | 0.001 | 0.509 | 0.926 |
|  | ADHD | DM | 671 | 0.010 | 0.000 | 0.926 | 0.926 |
|  | ADHD | WK | 750 | -0.051 | 0.000 | 0.639 | 0.926 |
|  | ADHD | FF | 365 | 0.021 | 0.000 | 0.884 | 0.926 |
|  | ASD × ADHD | TP | 606 | 0.725 | 0.015 | 0.003 | 0.010 |
|  | ASD × ADHD | DM | 671 | 0.115 | 0.000 | 0.604 | 0.805 |
|  | ASD × ADHD | WK | 750 | 0.274 | 0.002 | 0.186 | 0.372 |
|  | ASD × ADHD | FF | 365 | 0.046 | 0.000 | 0.886 | 0.886 |

Despite roughly halving the sample (n = 438–820 against 854–1455), estimates increased rather than attenuating. The autism main effect on gaze synchrony ranged from −0.85 to −1.15 SD (all q ≤ 0.001) and the ASD × ADHD interaction from +0.68 to +1.01 SD, reaching significance in all four contexts — including *Fun with Fractals*, the abstract non-social stimulus, where the interaction is not significant in the full sample (q = 0.018).

The direction of this change is informative. Random measurement heterogeneity attenuates standardized effects, so removing it should enlarge them, and that is what is observed; acquisition heterogeneity therefore suppressed these effects in the full sample rather than generating them. We treat this as corroborative rather than decisive, since a configuration-restricted subsample may differ from the full sample in composition as well as in measurement quality.

That the interaction reaches significance for *Fun with Fractals* under this restriction bears on the breadth of the effect. The attenuation associated with co-occurring ADHD is not confined to socially or narratively structured stimuli once measurement heterogeneity is removed.

**S3.2. Models without acquisition covariates**

Tracker configuration and screen shape are carried as covariates throughout because they are unbalanced across diagnostic groups (Table S2). To show what that adjustment contributes, we refitted the primary models adjusting for age and sex alone (Supplementary Table S8).

**Supplementary Table S8.** The primary 2 × 2 analysis of covariance refitted without tracker configuration and screen shape, adjusted for age and sex alone. Estimates differ from the primary specification by at most 0.04 SD for gaze metrics; the pupil interaction in *The Present* is the only term whose significance depends on the specification. Cell sizes per paradigm (TD / ASD only / ADHD only / comorbid): TP 150/60/858/189; DM 194/71/915/210; WK 198/78/963/216; FF 145/45/540/124.

| Metric | Term | Video | n | β | η_p_^2^ | p | q |
| --- | --- | --- | --- | --- | --- | --- | --- |
| Gaze  ISC  Magnitude | ASD | TP | 1257 | -0.759 | 0.020 | < 0.001 | < 0.001 |
|  | ASD | DM | 1390 | -0.753 | 0.021 | < 0.001 | < 0.001 |
|  | ASD | WK | 1455 | -0.600 | 0.014 | < 0.001 | < 0.001 |
|  | ASD | FF | 854 | -0.473 | 0.009 | 0.006 | 0.006 |
|  | ADHD | TP | 1257 | -0.102 | 0.001 | 0.243 | 0.243 |
|  | ADHD | DM | 1390 | -0.129 | 0.002 | 0.102 | 0.243 |
|  | ADHD | WK | 1455 | -0.092 | 0.001 | 0.237 | 0.243 |
|  | ADHD | FF | 854 | -0.131 | 0.002 | 0.161 | 0.243 |
|  | ASD × ADHD | TP | 1257 | 0.459 | 0.006 | 0.007 | 0.009 |
|  | ASD × ADHD | DM | 1390 | 0.607 | 0.011 | < 0.001 | < 0.001 |
|  | ASD × ADHD | WK | 1455 | 0.448 | 0.006 | 0.003 | 0.006 |
|  | ASD × ADHD | FF | 854 | 0.298 | 0.003 | 0.128 | 0.128 |
| Absolute  Gaze  Phase | ASD | TP | 1257 | -0.040 | 0.000 | 0.795 | 0.795 |
|  | ASD | DM | 1390 | 0.417 | 0.006 | 0.003 | 0.012 |
|  | ASD | WK | 1455 | 0.350 | 0.005 | 0.010 | 0.019 |
|  | ASD | FF | 854 | 0.073 | 0.000 | 0.673 | 0.795 |
|  | ADHD | TP | 1257 | 0.039 | 0.000 | 0.662 | 0.882 |
|  | ADHD | DM | 1390 | 0.068 | 0.001 | 0.389 | 0.779 |
|  | ADHD | WK | 1455 | 0.142 | 0.002 | 0.071 | 0.284 |
|  | ADHD | FF | 854 | -0.010 | 0.000 | 0.916 | 0.916 |
|  | ASD × ADHD | TP | 1257 | 0.094 | 0.000 | 0.586 | 0.654 |
|  | ASD × ADHD | DM | 1390 | -0.386 | 0.004 | 0.015 | 0.060 |
|  | ASD × ADHD | WK | 1455 | -0.240 | 0.002 | 0.118 | 0.235 |
|  | ASD × ADHD | FF | 854 | -0.089 | 0.000 | 0.654 | 0.654 |
| Pupil  ISC | ASD | TP | 757 | -0.766 | 0.020 | < 0.001 | < 0.001 |
|  | ASD | DM | 827 | -0.450 | 0.008 | 0.011 | 0.023 |
|  | ASD | WK | 917 | -0.337 | 0.004 | 0.047 | 0.062 |
|  | ASD | FF | 401 | -0.124 | 0.001 | 0.632 | 0.632 |
|  | ADHD | TP | 757 | -0.028 | 0.000 | 0.820 | 0.820 |
|  | ADHD | DM | 827 | 0.037 | 0.000 | 0.726 | 0.820 |
|  | ADHD | WK | 917 | -0.070 | 0.001 | 0.493 | 0.820 |
|  | ADHD | FF | 401 | 0.078 | 0.001 | 0.571 | 0.820 |
|  | ASD × ADHD | TP | 757 | 0.487 | 0.007 | 0.027 | 0.109 |
|  | ASD × ADHD | DM | 827 | 0.057 | 0.000 | 0.780 | 0.944 |
|  | ASD × ADHD | WK | 917 | 0.260 | 0.002 | 0.179 | 0.357 |
|  | ASD × ADHD | FF | 401 | 0.021 | 0.000 | 0.944 | 0.944 |

For gaze synchrony the adjustment makes almost no difference. The autism main effect moved by at most 0.03 SD in any context (unadjusted −0.76, −0.75, −0.60 and −0.47 against adjusted −0.78, −0.78, −0.60 and −0.45), and the interaction by at most 0.04 SD (+0.46, +0.61, +0.45, +0.30 against +0.50, +0.65, +0.46, +0.28). Every term significant in one specification was significant in the other, and the ADHD main effect remained null throughout (all q ≥ 0.24). The acquisition covariates are retained because the imbalance they address is real, not because they alter the conclusion.

The pupil interaction is the one place the specifications diverge. In *The Present* it is significant in the adjusted model (β = +0.56, q = 0.041) and not in the unadjusted one (β = +0.49, q = 0.109). That result is already treated as exploratory in the main text, and its dependence on specification is a further reason not to interpret it.

**S3.3. Adjustment for co-occurring psychopathology**

Children with co-occurring autism and ADHD carry more diagnoses and higher general psychopathology than children with autism alone (Section S6), raising the possibility that the interaction reflects those differences rather than the diagnostic combination. We repeated the models with oppositional or conduct diagnosis and CBCL Total T-score as additional covariates (Supplementary Table S9).

**Supplementary Table S9.** The 2 × 2 analysis of covariance additionally adjusted for oppositional or conduct diagnosis and CBCL Total T-score. Reported as a sensitivity analysis rather than as the primary model, since the CBCL partly indexes the conditions being contrasted. The interaction strengthens under this adjustment. Cell sizes per paradigm (TD / ASD only / ADHD only / comorbid): TP 150/60/858/189; DM 194/71/915/210; WK 198/78/963/216; FF 145/45/540/124.

| Metric | Term | Video | n | β | η_p_^2^ | p | q |
| --- | --- | --- | --- | --- | --- | --- | --- |
| Gaze  ISC  Magnitude | ASD | TP | 1257 | -0.881 | 0.025 | < 0.001 | < 0.001 |
|  | ASD | DM | 1390 | -0.831 | 0.023 | < 0.001 | < 0.001 |
|  | ASD | WK | 1455 | -0.653 | 0.015 | < 0.001 | < 0.001 |
|  | ASD | FF | 854 | -0.572 | 0.012 | 0.002 | 0.002 |
|  | ADHD | TP | 1257 | -0.155 | 0.002 | 0.105 | 0.140 |
|  | ADHD | DM | 1390 | -0.167 | 0.003 | 0.065 | 0.130 |
|  | ADHD | WK | 1455 | -0.076 | 0.001 | 0.388 | 0.388 |
|  | ADHD | FF | 854 | -0.229 | 0.006 | 0.032 | 0.129 |
|  | ASD × ADHD | TP | 1257 | 0.593 | 0.009 | 0.001 | 0.001 |
|  | ASD × ADHD | DM | 1390 | 0.698 | 0.013 | < 0.001 | < 0.001 |
|  | ASD × ADHD | WK | 1455 | 0.506 | 0.007 | 0.001 | 0.002 |
|  | ASD × ADHD | FF | 854 | 0.384 | 0.004 | 0.060 | 0.060 |
| Absolute  Gaze  Phase | ASD | TP | 1257 | -0.009 | 0.000 | 0.957 | 0.957 |
|  | ASD | DM | 1390 | 0.479 | 0.008 | 0.001 | 0.006 |
|  | ASD | WK | 1455 | 0.340 | 0.004 | 0.019 | 0.038 |
|  | ASD | FF | 854 | 0.047 | 0.000 | 0.802 | 0.957 |
|  | ADHD | TP | 1257 | 0.082 | 0.001 | 0.401 | 0.535 |
|  | ADHD | DM | 1390 | 0.124 | 0.001 | 0.175 | 0.437 |
|  | ADHD | WK | 1455 | 0.111 | 0.001 | 0.218 | 0.437 |
|  | ADHD | FF | 854 | -0.038 | 0.000 | 0.733 | 0.733 |
|  | ASD × ADHD | TP | 1257 | 0.038 | 0.000 | 0.832 | 0.832 |
|  | ASD × ADHD | DM | 1390 | -0.452 | 0.005 | 0.007 | 0.027 |
|  | ASD × ADHD | WK | 1455 | -0.242 | 0.002 | 0.134 | 0.269 |
|  | ASD × ADHD | FF | 854 | -0.076 | 0.000 | 0.718 | 0.832 |
| Pupil  ISC | ASD | TP | 757 | -0.803 | 0.022 | < 0.001 | < 0.001 |
|  | ASD | DM | 827 | -0.513 | 0.010 | 0.005 | 0.011 |
|  | ASD | WK | 917 | -0.409 | 0.006 | 0.022 | 0.029 |
|  | ASD | FF | 401 | -0.094 | 0.000 | 0.735 | 0.735 |
|  | ADHD | TP | 757 | -0.057 | 0.000 | 0.653 | 0.662 |
|  | ADHD | DM | 827 | -0.052 | 0.000 | 0.662 | 0.662 |
|  | ADHD | WK | 917 | -0.128 | 0.001 | 0.269 | 0.662 |
|  | ADHD | FF | 401 | 0.140 | 0.002 | 0.376 | 0.662 |
|  | ASD × ADHD | TP | 757 | 0.540 | 0.008 | 0.015 | 0.058 |
|  | ASD × ADHD | DM | 827 | 0.139 | 0.001 | 0.504 | 0.672 |
|  | ASD × ADHD | WK | 917 | 0.338 | 0.003 | 0.092 | 0.185 |
|  | ASD × ADHD | FF | 401 | 0.028 | 0.000 | 0.928 | 0.928 |

The interaction strengthened in every context (*The Present*: β = +0.59, q = 0.001; *Despicable Me*: β = +0.70, q < 0.001; *Diary of a Wimpy Kid*: β = +0.51, q = 0.002; *Fun with Fractals*: β = +0.38, q = 0.060), as did the autism main effects. General psychopathology therefore suppresses the observed pattern rather than producing it, which is the direction expected given that every clinical dimension distinguishing the two autism groups is elevated in the comorbid group (Section S6).

Two other terms changed status under this adjustment. The phase interaction during *Despicable Me* became significant (β = −0.45, q = 0.027), and the pupil interaction during *The Present* fell short of significance (β = +0.54, q = 0.058 against 0.041 in the primary model). Neither alters the interpretation: phase is not interpreted for the reasons given in Section S4, and the pupil interaction is treated as exploratory throughout.

This is reported as a sensitivity analysis rather than as the primary specification, because the CBCL partly indexes the conditions being contrasted; adjusting for it removes variance attributable to autism and ADHD themselves, and the resulting estimates should be read as a bound rather than as a better-specified model.

**S3.4. Robust regression**

Because the autism-without-ADHD cell is small, a few extreme observations within it could shift its mean. We refitted the models by Huber robust regression, which down-weights observations with large residuals (Supplementary Table S10).

**Supplementary Table S10.** Huber robust regression versions of the 2 × 2 models, down-weighting observations with large residuals, given the size of the autism-without-ADHD cell. Cell sizes per paradigm (TD / ASD only / ADHD only / comorbid): TP 150/60/858/189; DM 194/71/915/210; WK 198/78/963/216; FF 145/45/540/124.

| Metric | Term | Video | n | β | p | q |
| --- | --- | --- | --- | --- | --- | --- |
| Gaze  ISC  Magnitude | ASD | TP | 1257 | -0.717 | < 0.001 | < 0.001 |
|  | ASD | DM | 1390 | -0.636 | < 0.001 | < 0.001 |
|  | ASD | WK | 1455 | -0.497 | < 0.001 | < 0.001 |
|  | ASD | FF | 854 | -0.442 | 0.005 | 0.005 |
|  | ADHD | TP | 1257 | -0.097 | 0.193 | 0.257 |
|  | ADHD | DM | 1390 | -0.109 | 0.120 | 0.249 |
|  | ADHD | WK | 1455 | -0.062 | 0.315 | 0.315 |
|  | ADHD | FF | 854 | -0.133 | 0.125 | 0.249 |
|  | ASD × ADHD | TP | 1257 | 0.429 | 0.003 | 0.003 |
|  | ASD × ADHD | DM | 1390 | 0.481 | < 0.001 | 0.001 |
|  | ASD × ADHD | WK | 1455 | 0.398 | 0.001 | 0.001 |
|  | ASD × ADHD | FF | 854 | 0.243 | 0.176 | 0.176 |
| Absolute  Gaze  Phase | ASD | TP | 1257 | 0.033 | 0.523 | 0.523 |
|  | ASD | DM | 1390 | 0.167 | < 0.001 | 0.002 |
|  | ASD | WK | 1455 | 0.122 | 0.043 | 0.086 |
|  | ASD | FF | 854 | 0.108 | 0.142 | 0.189 |
|  | ADHD | TP | 1257 | -0.005 | 0.883 | 0.883 |
|  | ADHD | DM | 1390 | 0.040 | 0.154 | 0.307 |
|  | ADHD | WK | 1455 | 0.098 | 0.006 | 0.023 |
|  | ADHD | FF | 854 | -0.025 | 0.537 | 0.717 |
|  | ASD × ADHD | TP | 1257 | 0.022 | 0.703 | 0.703 |
|  | ASD × ADHD | DM | 1390 | -0.126 | 0.020 | 0.080 |
|  | ASD × ADHD | WK | 1455 | -0.107 | 0.116 | 0.232 |
|  | ASD × ADHD | FF | 854 | -0.042 | 0.621 | 0.703 |
| Pupil  ISC | ASD | TP | 757 | -0.742 | < 0.001 | < 0.001 |
|  | ASD | DM | 827 | -0.429 | 0.011 | 0.022 |
|  | ASD | WK | 917 | -0.327 | 0.027 | 0.035 |
|  | ASD | FF | 401 | -0.133 | 0.597 | 0.597 |
|  | ADHD | TP | 757 | -0.102 | 0.290 | 0.445 |
|  | ADHD | DM | 827 | 0.077 | 0.445 | 0.445 |
|  | ADHD | WK | 917 | -0.079 | 0.381 | 0.445 |
|  | ADHD | FF | 401 | 0.123 | 0.357 | 0.445 |
|  | ASD × ADHD | TP | 757 | 0.539 | 0.002 | 0.008 |
|  | ASD × ADHD | DM | 827 | 0.035 | 0.857 | 0.857 |
|  | ASD × ADHD | WK | 917 | 0.302 | 0.073 | 0.145 |
|  | ASD × ADHD | FF | 401 | 0.083 | 0.775 | 0.857 |

For gaze synchrony, estimates attenuated by 10–20% in most contexts and the inferential pattern was unchanged. The autism main effect remained significant in all four (β = −0.44 to −0.72, all q ≤ 0.005) and the interaction in the same three as the primary models (*The Present*: β = +0.43, q = 0.003; *Despicable Me*: β = +0.48, q = 0.001; *Diary of a Wimpy Kid*: β = +0.40, q = 0.001), with *Fun with Fractals* again short of significance (β = +0.24, q = 0.176).

Pupil ISC behaved similarly. The autism main effect was significant in three contexts (*The Present*: β = −0.74, q = 4.9 × 10⁻⁶; *Despicable Me*: β = −0.43, q = 0.022; *Diary of a Wimpy Kid*: β = −0.33, q = 0.035), and the interaction in *The Present* survived down-weighting with a smaller q than in the primary model (β = +0.54, q = 0.008).

Absolute gaze phase behaved differently, and the difference is informative. Its autism main effects attenuated by roughly 60% rather than 10–20% — from +0.41 to +0.17 in *Despicable Me* and from +0.35 to +0.12 in *Diary of a Wimpy Kid* — leaving only the *Despicable Me* effect significant among them (q = 0.002) and none of the interactions anywhere (all q ≥ 0.08). An ADHD main effect on phase during *Diary of a Wimpy Kid*, absent from the primary models, reached significance (β = +0.10, q = 0.023). The phase results are thus not merely a re-expression of magnitude (Section S4) but are also four times more sensitive to extreme observations than the magnitude results computed from the same recordings. Two independent properties argue against interpreting them.

**Supplementary Section S4: Absolute Gaze Phase**

The complex correlation yields two components, a modulus and an argument, and the argument might be expected to index spatial misalignment independently of the modulus. This section documents why that interpretation is not supported.

**S4.1. Phase and magnitude are not independent**

The argument of a complex correlation is estimated with precision that scales with its modulus: a participant whose gaze is weakly entrained to the normative trajectory will show a larger apparent angular offset for purely statistical reasons. Within each viewing context, absolute phase was substantially and negatively associated with magnitude (Supplementary Table S11): Pearson r = −0.399 (*The Present*), −0.437 (*Despicable Me*), −0.451 (*Diary of a Wimpy Kid*) and −0.319 (*Fun with Fractals*), the two metrics sharing 10–20% of their variance.

**Supplementary Table S11.** Association between absolute gaze phase and gaze ISC magnitude within each viewing context, as Pearson and Spearman correlations and the proportion of shared variance. The two metrics are components of the same complex correlation and are not independent.

| Video | n | Pearson’s r | Spearman’s ρ | R^2^ |
| --- | --- | --- | --- | --- |
| TP | 1775 | -0.399 | -0.226 | 0.159 |
| DM | 1959 | -0.437 | -0.236 | 0.191 |
| WK | 2042 | -0.451 | -0.248 | 0.203 |
| FF | 1186 | -0.319 | -0.200 | 0.102 |

The Spearman correlations are roughly half the Pearson values (ρ = −0.200 to −0.248), which indicates that the linear association is carried disproportionately by participants at the extremes of the distribution. This is the same property that makes the phase group effects fragile under robust regression (Section S3.4).

**S4.2. Observed phase against the floor implied by magnitude alone**

To quantify how much angular offset this dependence produces in the absence of any true rotation, we simulated for each video first-order autoregressive surrogate participants (12 moduli, 120 surrogates each) whose gaze related to the normative trajectory with a given modulus and the empirically measured gaze autocorrelation (lag-1 r = 0.976–0.985), but with no true angular offset, and interpolated the expected phase for each participant’s own magnitude. Group means are compared with their floors in Supplementary Table S12 and the full distributions in main text Figure 4.

**Supplementary Table S12.** Observed absolute gaze phase by diagnostic group against the phase expected from each participant’s own ISC magnitude under a null of no true angular offset, simulated using each video’s measured gaze autocorrelation. Mean mag is the group mean modulus, excess the observed mean minus the simulated floor. Negative values indicate group means below a floor that a true angular offset could not produce, and reflect a small upward bias in the parametric surrogate.

| Video | Diagnosis | n | Mean mag | Observed phase | Floor phase | excess |
| --- | --- | --- | --- | --- | --- | --- |
| TP | TD | 147 | 0.678 | 0.077 | 0.067 | 0.010 |
|  | ADHD only | 850 | 0.668 | 0.082 | 0.069 | 0.013 |
|  | ASD only | 59 | 0.618 | 0.075 | 0.081 | -0.006 |
|  | ASD and ADHD | 188 | 0.643 | 0.097 | 0.075 | 0.022 |
|  | Other | 513 | 0.671 | 0.075 | 0.068 | 0.007 |
| DM | TD | 194 | 0.614 | 0.083 | 0.079 | 0.004 |
|  | ADHD only | 902 | 0.605 | 0.082 | 0.082 | 0.001 |
|  | ASD only | 71 | 0.552 | 0.139 | 0.097 | 0.042 |
|  | ASD and ADHD | 209 | 0.594 | 0.086 | 0.084 | 0.002 |
|  | Other | 563 | 0.604 | 0.088 | 0.082 | 0.006 |
| WK | TD | 196 | 0.609 | 0.069 | 0.087 | -0.018 |
|  | ADHD only | 954 | 0.607 | 0.081 | 0.088 | -0.007 |
|  | ASD only | 77 | 0.571 | 0.086 | 0.099 | -0.013 |
|  | ASD and ADHD | 215 | 0.596 | 0.098 | 0.091 | 0.007 |
|  | Other | 578 | 0.604 | 0.079 | 0.089 | -0.010 |
| FF | TD | 144 | 0.520 | 0.099 | 0.112 | -0.013 |
|  | ADHD only | 530 | 0.507 | 0.101 | 0.116 | -0.014 |
|  | ASD only | 44 | 0.491 | 0.115 | 0.121 | -0.007 |
|  | ASD and ADHD | 124 | 0.494 | 0.101 | 0.121 | -0.020 |
|  | Other | 332 | 0.503 | 0.107 | 0.118 | -0.011 |

One group mean departed appreciably from its floor: autism without ADHD during *Despicable Me*, observed 0.139 against an expected 0.097, an excess of 0.042 radians. Every other group mean in every context lay between −0.020 and +0.022 radians of its floor. The excess for that one cell is also the source of the larger of the two apparent phase main effects in the primary models, and that cell contains 71 participants.

Several group means fell below their floors — all five groups during *Fun with Fractals* and four of five during *Diary of a Wimpy Kid*. A true angular offset cannot be negative, so this indicates that the parametric surrogate slightly overestimates the floor in absolute terms. The formal tests below therefore rely on within-video contrasts, in which a uniform offset cancels, and on a non-parametric alternative requiring no simulation at all.

**S4.3. Group models on magnitude-adjusted phase**

We refitted the 2 × 2 models to phase adjusted for magnitude by two independent routes. Subtracting the simulated floor (Supplementary Table S13) left no main effect or interaction significant in any context, the smallest corrected q being 0.064 for the autism main effect during *Despicable Me*. Residualizing phase on a cubic B-spline (five degrees of freedom) in magnitude within each video, which assumes nothing about the data-generating process (Supplementary Table S14), likewise left nothing significant, the smallest q being 0.144 for the autism main effect during *The Present*.

**Supplementary Table S13.** The 2 × 2 analysis of covariance on absolute gaze phase after subtracting the magnitude-implied floor of Table S12. Sample sizes are slightly smaller than in the primary models because the floor is undefined for participants whose magnitude falls outside the simulated grid. Cell sizes per paradigm (TD / ASD only / ADHD only / comorbid): TP 147/59/850/188; DM 194/71/902/209; WK 196/77/954/215; FF 144/44/530/124.

| Term | Video | n | β | η_p_^2^ | p | q |
| --- | --- | --- | --- | --- | --- | --- |
| ASD | TP | 1244 | -0.113 | 0.000 | 0.467 | 0.698 |
|  | DM | 1376 | 0.339 | 0.004 | 0.016 | 0.064 |
|  | WK | 1442 | 0.087 | 0.000 | 0.524 | 0.698 |
|  | FF | 842 | 0.065 | 0.000 | 0.710 | 0.710 |
| ADHD | TP | 1244 | 0.033 | 0.000 | 0.713 | 0.951 |
|  | DM | 1376 | -0.028 | 0.000 | 0.730 | 0.951 |
|  | WK | 1442 | 0.142 | 0.002 | 0.079 | 0.315 |
|  | FF | 842 | -0.006 | 0.000 | 0.951 | 0.951 |
| ASD × ADHD | TP | 1244 | 0.150 | 0.001 | 0.392 | 0.710 |
|  | DM | 1376 | -0.325 | 0.003 | 0.041 | 0.166 |
|  | WK | 1442 | 0.058 | 0.000 | 0.710 | 0.710 |
|  | FF | 842 | -0.112 | 0.000 | 0.577 | 0.710 |

**Supplementary Table S14.** As Table S13, but with phase residualized non-parametrically on a cubic B-spline in magnitude within each video, requiring no simulation assumption.

| Term | Video | n | β | η_p_^2^ | p | q |
| --- | --- | --- | --- | --- | --- | --- |
| ASD | TP | 1257 | -0.323 | 0.004 | 0.036 | 0.144 |
|  | DM | 1390 | 0.127 | 0.001 | 0.366 | 0.488 |
|  | WK | 1455 | 0.154 | 0.001 | 0.258 | 0.488 |
|  | FF | 854 | -0.050 | 0.000 | 0.774 | 0.774 |
| ADHD | TP | 1257 | 0.030 | 0.000 | 0.740 | 0.740 |
|  | DM | 1390 | -0.034 | 0.000 | 0.673 | 0.740 |
|  | WK | 1455 | 0.122 | 0.002 | 0.129 | 0.515 |
|  | FF | 854 | -0.038 | 0.000 | 0.696 | 0.740 |
| ASD × ADHD | TP | 1257 | 0.318 | 0.003 | 0.067 | 0.266 |
|  | DM | 1390 | -0.124 | 0.000 | 0.437 | 0.783 |
|  | WK | 1455 | -0.084 | 0.000 | 0.587 | 0.783 |
|  | FF | 854 | 0.028 | 0.000 | 0.888 | 0.888 |

**Supplementary Figure S4.** Absolute gaze phase |θ| by ASD and ADHD status. One panel per viewing context. Autism status is encoded by color and ADHD status by horizontal position, matching main text Figure 3, of which this is the row omitted from the main text. Points are individual participants; boxes give the median and interquartile range; markers and vertical bars give the cell mean and its 95% confidence interval. Point opacity, size and jitter width are scaled by cell size, and cell *n* is printed along the bottom of each panel. The heterogeneous “Other” diagnostic group is excluded. The group differences visible here are not interpreted: Section S4 shows that they are a re-expression of the corresponding differences in synchrony magnitude, and that they do not survive adjustment for the phase each participant’s own magnitude implies (Tables S13 and S14).

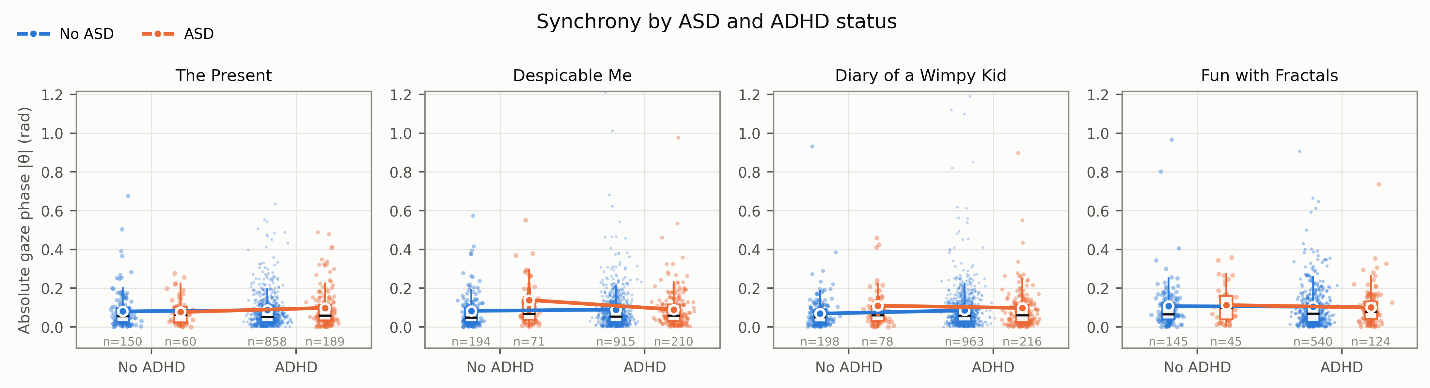

The two approaches disagree about which context comes closest to significance — the simulation favoring *Despicable Me*, the spline favoring *The Present* — and the signs of several terms differ between them. Disagreement of this kind is what is expected when no stable signal is present. The two analyses also run on slightly different samples (n = 842–1442 against 854–1455), since the simulated floor is undefined for participants whose magnitude falls outside the simulated grid.

**S4.4. Generality**

We report this as a negative methodological result rather than as a feature of these data. Any application of complex-correlation or phase-based synchrony to smooth, strongly autocorrelated signals faces the same confound, and gaze position is strongly autocorrelated by construction. A magnitude-matched null is necessary before a phase difference between groups can be interpreted.

**Supplementary Section S5: Coarse Oculomotor Measures**

**S5.1. What these measures index**

To establish what the inter-subject framework adds over stimulus-independent summaries of eye movement, and to test the mechanism proposed for the interaction, we computed ten non-inter-subject measures from the same 30 Hz output grid: gaze-shift rate at 30 and 60 °/s thresholds, mean dwell duration between shifts, proportion of time shifting, mean scan-path speed, bivariate contour ellipse area (BCEA), horizontal and vertical positional standard deviation, overall dispersion, and proportion of time within the central third of the screen in both dimensions. Mean dwell duration is valid viewing time divided by the number of shifts, and is therefore the exact per-subject reciprocal of gaze-shift rate.

Two constraints follow from the sampling rate. A saccade lasts 30–80 ms, one to two samples at 30 Hz, and the common 10 Hz low-pass attenuates peak velocities; these measures therefore index resolvable gaze shifts rather than saccades, and dwell segments rather than fixations in the oculomotor sense. Absolute values are consequently not comparable with published norms. They are comparable across participants, since every recording passed through an identical pipeline at an identical final rate, which is what the group comparison requires.

**S5.2. Group models**

The 2 × 2 models applied to the ISC metrics were applied unchanged to each measure (Supplementary Table S15). Across 120 tests — ten measures, four contexts, three model terms — two survived correction, both for mean dwell duration during *Diary of a Wimpy Kid*: an autism main effect (β = +0.51, q = 0.0003) and an ASD × ADHD interaction (β = −0.47, q = 0.007). No other term reached significance in any context, the smallest remaining q being 0.118. The measures most directly comparable with standard oculomotor reporting — gaze-shift rate, dispersion, bivariate contour ellipse area and scan-path speed — were undifferentiated by diagnostic group throughout.

**Supplementary Table S15.** The 2 × 2 analysis of covariance applied to ten coarse, stimulus-independent gaze measures computed from the same 30 Hz grid, adjusted for age, sex, tracker configuration and screen shape. Benjamini-Hochberg q within each measure family across the four contexts. Measured at 30 Hz after a common 10 Hz low-pass, these index resolvable gaze shifts rather than saccades.

| Measure | Term | Video | n | β | η_p_^2^ | p | q |
| --- | --- | --- | --- | --- | --- | --- | --- |
| Gaze-shift rate  >30°/s (shifts/s) | ASD | TP | 1257 | -0.010 | 0.000 | 0.942 | 0.961 |
|  | ASD | DM | 1390 | -0.058 | 0.000 | 0.645 | 0.961 |
|  | ASD | WK | 1455 | -0.104 | 0.001 | 0.399 | 0.961 |
|  | ASD | FF | 854 | -0.008 | 0.000 | 0.961 | 0.961 |
|  | ADHD | TP | 1257 | 0.109 | 0.002 | 0.176 | 0.234 |
|  | ADHD | DM | 1390 | -0.008 | 0.000 | 0.908 | 0.908 |
|  | ADHD | WK | 1455 | 0.104 | 0.001 | 0.155 | 0.234 |
|  | ADHD | FF | 854 | 0.120 | 0.002 | 0.164 | 0.234 |
|  | ASD × ADHD | TP | 1257 | 0.056 | 0.000 | 0.718 | 0.718 |
|  | ASD × ADHD | DM | 1390 | 0.089 | 0.000 | 0.534 | 0.712 |
|  | ASD × ADHD | WK | 1455 | 0.123 | 0.001 | 0.383 | 0.712 |
|  | ASD × ADHD | FF | 854 | 0.220 | 0.002 | 0.219 | 0.712 |
| Gaze-shift rate  >60°/s (shifts/s) | ASD | TP | 1257 | 0.015 | 0.000 | 0.908 | 0.908 |
|  | ASD | DM | 1390 | -0.114 | 0.001 | 0.369 | 0.739 |
|  | ASD | WK | 1455 | -0.148 | 0.001 | 0.224 | 0.739 |
|  | ASD | FF | 854 | 0.057 | 0.000 | 0.714 | 0.908 |
|  | ADHD | TP | 1257 | 0.120 | 0.002 | 0.107 | 0.239 |
|  | ADHD | DM | 1390 | -0.098 | 0.001 | 0.179 | 0.239 |
|  | ADHD | WK | 1455 | 0.050 | 0.000 | 0.485 | 0.485 |
|  | ADHD | FF | 854 | 0.117 | 0.002 | 0.173 | 0.239 |
|  | ASD × ADHD | TP | 1257 | 0.058 | 0.000 | 0.685 | 0.685 |
|  | ASD × ADHD | DM | 1390 | 0.136 | 0.001 | 0.344 | 0.458 |
|  | ASD × ADHD | WK | 1455 | 0.146 | 0.001 | 0.291 | 0.458 |
|  | ASD × ADHD | FF | 854 | 0.212 | 0.002 | 0.231 | 0.458 |
| Mean dwell duration  between > 30^o^/s shifts (ms) | ASD | TP | 1257 | 0.095 | 0.000 | 0.492 | 0.657 |
|  | ASD | DM | 1390 | 0.138 | 0.001 | 0.279 | 0.559 |
|  | ASD | WK | 1455 | 0.515 | 0.011 | < 0.001 | < 0.001 |
|  | ASD | FF | 854 | 0.070 | 0.000 | 0.657 | 0.657 |
|  | ADHD | TP | 1257 | -0.045 | 0.000 | 0.575 | 0.921 |
|  | ADHD | DM | 1390 | -0.009 | 0.000 | 0.908 | 0.921 |
|  | ADHD | WK | 1455 | -0.008 | 0.000 | 0.921 | 0.921 |
|  | ADHD | FF | 854 | -0.053 | 0.000 | 0.542 | 0.921 |
|  | ASD × ADHD | TP | 1257 | -0.108 | 0.000 | 0.484 | 0.783 |
|  | ASD × ADHD | DM | 1390 | -0.040 | 0.000 | 0.783 | 0.783 |
|  | ASD × ADHD | WK | 1455 | -0.465 | 0.007 | 0.002 | 0.007 |
|  | ASD × ADHD | FF | 854 | -0.084 | 0.000 | 0.644 | 0.783 |
| Time shifting  >30^o^/s (%) | ASD | TP | 1257 | 0.006 | 0.000 | 0.963 | 0.963 |
|  | ASD | DM | 1390 | -0.074 | 0.000 | 0.559 | 0.963 |
|  | ASD | WK | 1455 | -0.088 | 0.000 | 0.470 | 0.963 |
|  | ASD | FF | 854 | 0.021 | 0.000 | 0.889 | 0.963 |
|  | ADHD | TP | 1257 | 0.095 | 0.001 | 0.226 | 0.350 |
|  | ADHD | DM | 1390 | -0.060 | 0.001 | 0.411 | 0.411 |
|  | ADHD | WK | 1455 | 0.082 | 0.001 | 0.253 | 0.350 |
|  | ADHD | FF | 854 | 0.095 | 0.002 | 0.263 | 0.350 |
|  | ASD × ADHD | TP | 1257 | 0.081 | 0.000 | 0.591 | 0.591 |
|  | ASD × ADHD | DM | 1390 | 0.121 | 0.001 | 0.398 | 0.547 |
|  | ASD × ADHD | WK | 1455 | 0.114 | 0.001 | 0.410 | 0.547 |
|  | ASD × ADHD | FF | 854 | 0.252 | 0.002 | 0.152 | 0.547 |
| Scan-path  speed (^o^/s) | ASD | TP | 1257 | 0.082 | 0.000 | 0.524 | 0.810 |
|  | ASD | DM | 1390 | -0.028 | 0.000 | 0.810 | 0.810 |
|  | ASD | WK | 1455 | 0.031 | 0.000 | 0.789 | 0.810 |
|  | ASD | FF | 854 | 0.085 | 0.000 | 0.556 | 0.810 |
|  | ADHD | TP | 1257 | 0.123 | 0.002 | 0.104 | 0.155 |
|  | ADHD | DM | 1390 | -0.022 | 0.000 | 0.746 | 0.746 |
|  | ADHD | WK | 1455 | 0.126 | 0.002 | 0.063 | 0.155 |
|  | ADHD | FF | 854 | 0.125 | 0.003 | 0.116 | 0.155 |
|  | ASD × ADHD | TP | 1257 | -0.017 | 0.000 | 0.908 | 0.908 |
|  | ASD × ADHD | DM | 1390 | 0.044 | 0.000 | 0.740 | 0.908 |
|  | ASD × ADHD | WK | 1455 | -0.025 | 0.000 | 0.848 | 0.908 |
|  | ASD × ADHD | FF | 854 | 0.098 | 0.000 | 0.553 | 0.908 |
| Bivariate contour  ellipse area  68% (deg²) | ASD | TP | 1257 | -0.097 | 0.001 | 0.361 | 0.613 |
|  | ASD | DM | 1390 | -0.059 | 0.000 | 0.613 | 0.613 |
|  | ASD | WK | 1455 | 0.151 | 0.001 | 0.183 | 0.613 |
|  | ASD | FF | 854 | 0.080 | 0.001 | 0.529 | 0.613 |
|  | ADHD | TP | 1257 | 0.032 | 0.000 | 0.599 | 0.797 |
|  | ADHD | DM | 1390 | -0.017 | 0.000 | 0.797 | 0.797 |
|  | ADHD | WK | 1455 | 0.112 | 0.002 | 0.096 | 0.385 |
|  | ADHD | FF | 854 | 0.055 | 0.001 | 0.431 | 0.797 |
|  | ASD × ADHD | TP | 1257 | 0.151 | 0.001 | 0.204 | 0.765 |
|  | ASD × ADHD | DM | 1390 | 0.039 | 0.000 | 0.765 | 0.765 |
|  | ASD × ADHD | WK | 1455 | -0.102 | 0.000 | 0.431 | 0.765 |
|  | ASD × ADHD | FF | 854 | -0.049 | 0.000 | 0.739 | 0.765 |
| Positional dispersion  (normalized units) | ASD | TP | 1257 | -0.169 | 0.002 | 0.171 | 0.498 |
|  | ASD | DM | 1390 | 0.000 | 0.000 | 1.000 | 1.000 |
|  | ASD | WK | 1455 | 0.140 | 0.001 | 0.275 | 0.498 |
|  | ASD | FF | 854 | 0.145 | 0.001 | 0.373 | 0.498 |
|  | ADHD | TP | 1257 | -0.024 | 0.000 | 0.735 | 0.735 |
|  | ADHD | DM | 1390 | -0.049 | 0.000 | 0.530 | 0.707 |
|  | ADHD | WK | 1455 | 0.161 | 0.003 | 0.034 | 0.137 |
|  | ADHD | FF | 854 | 0.067 | 0.001 | 0.458 | 0.707 |
|  | ASD × ADHD | TP | 1257 | 0.243 | 0.003 | 0.080 | 0.320 |
|  | ASD × ADHD | DM | 1390 | -0.055 | 0.000 | 0.720 | 0.728 |
|  | ASD × ADHD | WK | 1455 | -0.099 | 0.000 | 0.496 | 0.728 |
|  | ASD × ADHD | FF | 854 | -0.065 | 0.000 | 0.728 | 0.728 |
| Horizontal  position SD (^o^) | ASD | TP | 1257 | 0.000 | 0.000 | 0.999 | 0.999 |
|  | ASD | DM | 1390 | 0.126 | 0.001 | 0.293 | 0.586 |
|  | ASD | WK | 1455 | 0.139 | 0.001 | 0.231 | 0.586 |
|  | ASD | FF | 854 | 0.062 | 0.000 | 0.616 | 0.821 |
|  | ADHD | TP | 1257 | 0.006 | 0.000 | 0.921 | 0.921 |
|  | ADHD | DM | 1390 | 0.042 | 0.000 | 0.547 | 0.921 |
|  | ADHD | WK | 1455 | 0.123 | 0.002 | 0.075 | 0.299 |
|  | ADHD | FF | 854 | 0.011 | 0.000 | 0.875 | 0.921 |
|  | ASD × ADHD | TP | 1257 | 0.059 | 0.000 | 0.599 | 0.798 |
|  | ASD × ADHD | DM | 1390 | -0.191 | 0.001 | 0.159 | 0.635 |
|  | ASD × ADHD | WK | 1455 | -0.115 | 0.001 | 0.387 | 0.775 |
|  | ASD × ADHD | FF | 854 | -0.027 | 0.000 | 0.850 | 0.850 |
| Vertical  position SD (^o^) | ASD | TP | 1257 | -0.112 | 0.001 | 0.369 | 0.739 |
|  | ASD | DM | 1390 | -0.046 | 0.000 | 0.683 | 0.804 |
|  | ASD | WK | 1455 | 0.134 | 0.001 | 0.237 | 0.739 |
|  | ASD | FF | 854 | 0.035 | 0.000 | 0.804 | 0.804 |
|  | ADHD | TP | 1257 | 0.059 | 0.001 | 0.418 | 0.418 |
|  | ADHD | DM | 1390 | -0.074 | 0.001 | 0.258 | 0.361 |
|  | ADHD | WK | 1455 | 0.107 | 0.002 | 0.109 | 0.361 |
|  | ADHD | FF | 854 | 0.085 | 0.001 | 0.271 | 0.361 |
|  | ASD × ADHD | TP | 1257 | 0.158 | 0.001 | 0.258 | 0.689 |
|  | ASD × ADHD | DM | 1390 | 0.053 | 0.000 | 0.681 | 0.908 |
|  | ASD × ADHD | WK | 1455 | -0.122 | 0.001 | 0.345 | 0.689 |
|  | ASD × ADHD | FF | 854 | -0.012 | 0.000 | 0.941 | 0.941 |
| Time in central  third of display (%) | ASD | TP | 1257 | 0.026 | 0.000 | 0.852 | 0.883 |
|  | ASD | DM | 1390 | -0.129 | 0.001 | 0.351 | 0.701 |
|  | ASD | WK | 1455 | -0.232 | 0.002 | 0.075 | 0.298 |
|  | ASD | FF | 854 | -0.025 | 0.000 | 0.883 | 0.883 |
|  | ADHD | TP | 1257 | -0.009 | 0.000 | 0.910 | 0.910 |
|  | ADHD | DM | 1390 | 0.038 | 0.000 | 0.635 | 0.846 |
|  | ADHD | WK | 1455 | -0.168 | 0.003 | 0.030 | 0.118 |
|  | ADHD | FF | 854 | -0.051 | 0.000 | 0.584 | 0.846 |
|  | ASD × ADHD | TP | 1257 | -0.137 | 0.001 | 0.375 | 0.751 |
|  | ASD × ADHD | DM | 1390 | 0.047 | 0.000 | 0.764 | 0.764 |
|  | ASD × ADHD | WK | 1455 | 0.139 | 0.001 | 0.346 | 0.751 |
|  | ASD × ADHD | FF | 854 | -0.084 | 0.000 | 0.666 | 0.764 |

**S5.3. The dwell result is transformation-dependent**

Mean dwell duration and gaze-shift rate are exact per-subject reciprocals by construction. That one yields the only significant result in Table S15 while the other is null in the same context and metric family (autism main effect: β = −0.10, q = 0.96; interaction: β = +0.12, q = 0.71) indicates that the effect is not a difference in central tendency but a difference in the upper tail of the dwell distribution.

Supplementary Table S16 makes the asymmetry explicit. For the isolated-autism group during *Diary of a Wimpy Kid*, mean dwell duration is 1001 ms while the reciprocal of the group’s mean shift rate is 826 ms, a discrepancy of 175 ms produced by right skew; for the comorbid group the corresponding discrepancy is 78 ms. Standardizing a variable with this much skew gives extreme observations substantial leverage, and the reciprocal scale does not. We therefore report the dwell result but do not treat it as establishing a group difference in gaze-shift behavior, and we note that it concerns one measure in one of four contexts.

**Supplementary Table S16.** Comorbid minus isolated-autism contrasts on each coarse measure, with group means, Cohen’s d and whether the comorbid mean is the higher of the two. Mean dwell duration and shift rate are per-subject reciprocals, so their directions are necessarily opposed.

| Measure | Video | Mean_ASD_only_ | n_ASD_only_ | Mean_comorbid_ | n_comorbid_ | Group difference | Cohen’s d | Comorbid higher |
| --- | --- | --- | --- | --- | --- | --- | --- | --- |
| Gaze-shift rate  >30°/s (shifts/s) | TP | 1.05 | 60 | 1.14 | 189 | 0.091 | 0.168 | Yes |
|  | DM | 1.07 | 71 | 1.13 | 210 | 0.059 | 0.104 | Yes |
|  | WK | 1.21 | 78 | 1.34 | 216 | 0.133 | 0.286 | Yes |
|  | FF | 1.14 | 45 | 1.30 | 124 | 0.159 | 0.272 | Yes |
| Gaze-shift rate  >60°/s (shifts/s) | TP | 0.55 | 60 | 0.60 | 189 | 0.056 | 0.180 | Yes |
|  | DM | 0.59 | 71 | 0.61 | 210 | 0.024 | 0.081 | Yes |
|  | WK | 0.71 | 78 | 0.79 | 216 | 0.085 | 0.260 | Yes |
|  | FF | 0.60 | 45 | 0.70 | 124 | 0.100 | 0.239 | Yes |
| Mean dwell duration  between > 30^o^/s shifts (ms) | TP | 1066.95 | 60 | 995.99 | 189 | -70.962 | -0.216 | No |
|  | DM | 1056.45 | 71 | 1024.87 | 210 | -31.574 | -0.085 | No |
|  | WK | 1000.64 | 78 | 823.25 | 216 | -177.392 | -0.223 | No |
|  | FF | 962.63 | 45 | 913.75 | 124 | -48.881 | -0.130 | No |
| Time shifting  >30^o^/s (%) | TP | 7.15 | 60 | 7.89 | 189 | 0.741 | 0.179 | Yes |
|  | DM | 7.45 | 71 | 7.86 | 210 | 0.410 | 0.094 | Yes |
|  | WK | 8.49 | 78 | 9.53 | 216 | 1.036 | 0.253 | Yes |
|  | FF | 7.99 | 45 | 9.37 | 124 | 1.380 | 0.251 | Yes |
| Scan-path  speed (^o^/s) | TP | 9.51 | 60 | 10.06 | 189 | 0.552 | 0.161 | Yes |
|  | DM | 9.61 | 71 | 9.87 | 210 | 0.266 | 0.072 | Yes |
|  | WK | 10.58 | 78 | 11.34 | 216 | 0.756 | 0.191 | Yes |
|  | FF | 10.58 | 45 | 11.32 | 124 | 0.746 | 0.171 | Yes |
| Bivariate contour  ellipse area  68% (deg²) | TP | 47.42 | 60 | 51.56 | 189 | 4.144 | 0.179 | Yes |
|  | DM | 42.96 | 71 | 44.09 | 210 | 1.136 | 0.057 | Yes |
|  | WK | 38.15 | 78 | 39.95 | 216 | 1.808 | 0.097 | Yes |
|  | FF | 56.00 | 45 | 56.85 | 124 | 0.852 | 0.040 | Yes |
| Positional dispersion  (normalized units) | TP | 0.15 | 60 | 0.16 | 189 | 0.006 | 0.174 | Yes |
|  | DM | 0.15 | 71 | 0.15 | 210 | -0.002 | -0.072 | No |
|  | WK | 0.14 | 78 | 0.14 | 216 | 0.003 | 0.127 | Yes |
|  | FF | 0.16 | 45 | 0.16 | 124 | 0.002 | 0.082 | Yes |
| Horizontal  position SD (^o^) | TP | 3.29 | 60 | 3.32 | 189 | 0.026 | 0.030 | Yes |
|  | DM | 3.26 | 71 | 3.17 | 210 | -0.087 | -0.139 | No |
|  | WK | 2.95 | 78 | 3.01 | 216 | 0.058 | 0.078 | Yes |
|  | FF | 3.28 | 45 | 3.25 | 124 | -0.031 | -0.045 | No |
| Vertical  position SD (^o^) | TP | 2.02 | 60 | 2.16 | 189 | 0.139 | 0.270 | Yes |
|  | DM | 1.92 | 71 | 1.93 | 210 | 0.016 | 0.030 | Yes |
|  | WK | 1.80 | 78 | 1.84 | 216 | 0.039 | 0.087 | Yes |
|  | FF | 2.36 | 45 | 2.45 | 124 | 0.086 | 0.159 | Yes |
| Time in central  third of display (%) | TP | 72.54 | 60 | 70.68 | 189 | -1.863 | -0.110 | No |
|  | DM | 73.49 | 71 | 74.09 | 210 | 0.606 | 0.056 | Yes |
|  | WK | 77.31 | 78 | 76.13 | 216 | -1.182 | -0.080 | No |
|  | FF | 69.57 | 45 | 67.39 | 124 | -2.180 | -0.196 | No |

**S5.4. Directional test of the proposed mechanism**

The account under which more frequent gaze shifting disrupts prolonged idiosyncratic fixation makes an ordered prediction that these measures can falsify: comorbid children should shift gaze more often, and dwell less long, than children with autism alone. Table S16 gives the contrast for every measure.

The direction is consistent. Comorbid children showed higher gaze-shift rates at both velocity thresholds, a greater proportion of time shifting, faster scan paths, larger contour ellipse areas, greater vertical positional variability and shorter dwell durations than children with autism alone, in all four viewing contexts without exception. Effect sizes were uniformly small (|d| = 0.03 to 0.29). The three measures not showing this consistency were overall dispersion, horizontal positional standard deviation and proportion of time central.

Two reasons not to read this as confirmation. These measures are not independent — shift rate, proportion of time shifting, scan-path speed and dwell duration are near-equivalent summaries of the same quantity — so the consistency reflects perhaps two or three distinct constructs rather than seven. And none of the corresponding model terms in Table S15 approaches significance (shift-rate interaction, all q ≥ 0.46). The mechanism is a hypothesis these data fail to refute rather than one they support, and testing it properly requires oculomotor sampling finer than the 30 Hz common grid used here.

**S5.5. Scope of the result**

The group structure detected by inter-subject synchrony in these recordings was not recoverable from these coarse summaries of the same gaze data. The measures describe how the eyes move irrespective of what is on the screen, so the result locates the group differences in where gaze is directed relative to the stimulus, not in the gross dynamics of eye movement. It does not bear on content-referenced approaches such as region-of-interest or saliency-based measures, which were not computed (main text Section 4.5).

**Supplementary Figure S5.** Comorbid minus autism-without-ADHD differences on ten coarse, stimulus-independent gaze measures, expressed as Cohen’s *d*. Each video is shown by its own color and marker: *The Present* (circle), *Despicable Me* (square), *Diary of a Wimpy Kid* (diamond) and *Fun with Fractals* (triangle). Measures are ordered by their mean across videos. The account under which more frequent gaze shifting disrupts prolonged idiosyncratic fixation predicts positive values. The direction is consistent across the shifting-related measures in all four videos, but every estimate is small (|*d*| ≤ 0.29) and none of the corresponding terms in the 2 × 2 models reaches significance (Table S15). Mean dwell duration is the per-subject reciprocal of gaze-shift rate, so its sign is necessarily opposed to the others.

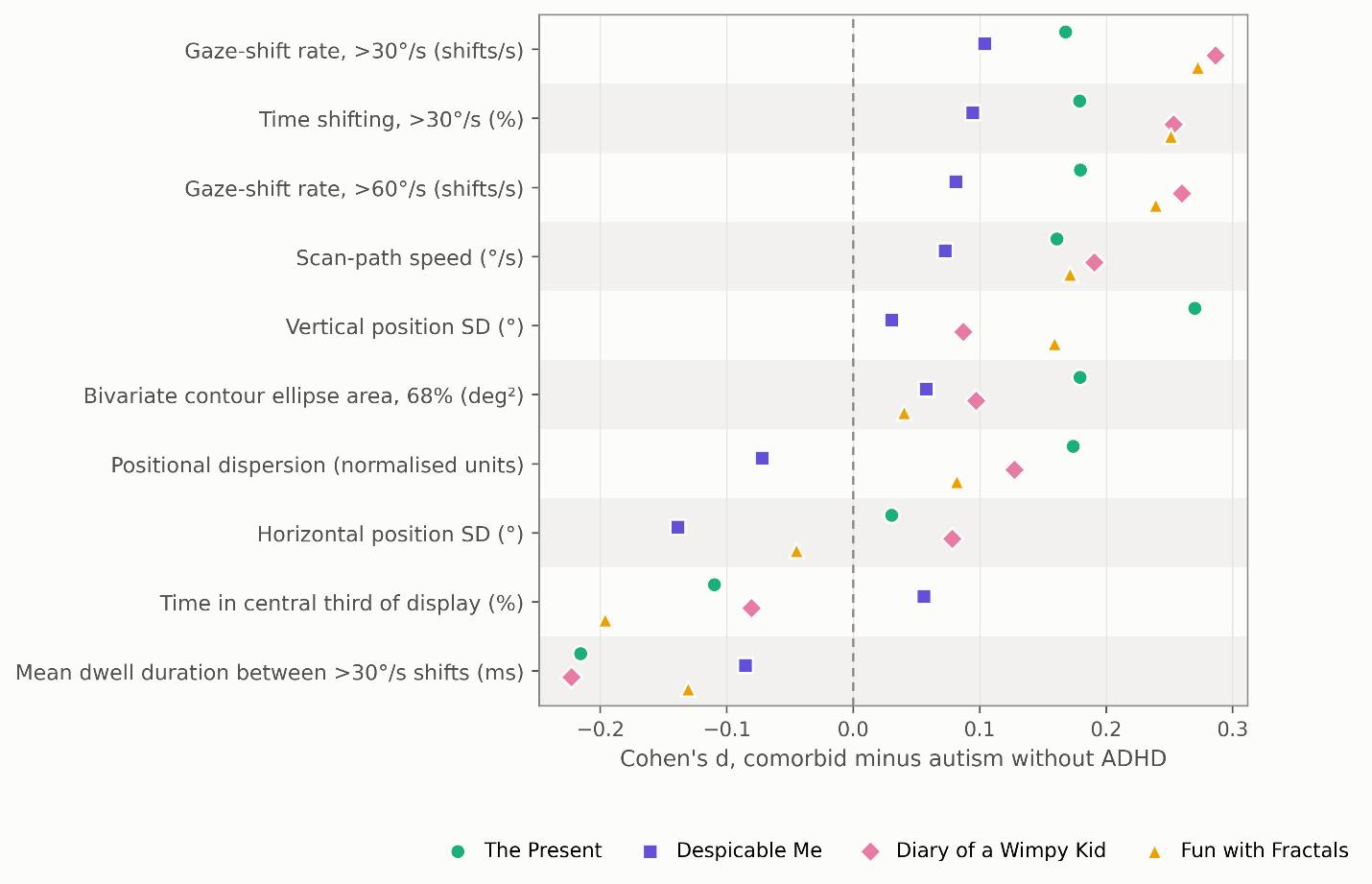

**Supplementary Section S6: Severity and Ascertainment**

**S6.1. The question**

The interaction reported in the main text could arise without any contribution from ADHD if the autism-without-ADHD group simply had more severe autism than the comorbid group. Children receiving an autism diagnosis without a co-occurring attentional diagnosis in a help-seeking cohort are an unusual subgroup, and it is plausible that they represent more prototypical or more severe presentations. We therefore compared the two autism groups on autistic trait severity, cognitive ability, general psychopathology and diagnostic load (Supplementary Table S17).

**Supplementary Table S17.** Comparison of the autism-without-ADHD and comorbid groups on autistic trait severity, its subscales, cognitive ability, general psychopathology, and diagnostic load. Welch’s t-test accommodates unequal variances; Mann-Whitney p values are given alongside for variables whose distributions are skewed. Cohen’s *d* is signed as comorbid minus autism-without-ADHD. Sample sizes vary across rows with instrument availability: SRS measures were available for 93 and 264 participants, Wechsler full-scale IQ for 63 and 219. SWAN Total is reported for completeness but is definitional, since it is the basis on which the groups are constituted.

| Variable | n_ASD_only_ | n_comorbid_ | Mean_ASD_only_ | Mean_comorbid_ | Welch t | p (Welch) | p (Mann-Whitney) | Cohen’s d |
| --- | --- | --- | --- | --- | --- | --- | --- | --- |
| SRS-2 Total T | 93 | 264 | 66.850 | 70.129 | -2.580 | 0.011 | 0.014 | 0.306 |
| Social Awareness | 93 | 264 | 62.011 | 66.735 | -3.465 | 0.001 | < 0.001 | 0.424 |
| Social Cognition | 93 | 264 | 65.280 | 67.580 | -1.594 | 0.113 | 0.150 | 0.191 |
| Social Communication | 93 | 264 | 67.194 | 69.564 | -1.895 | 0.060 | 0.078 | 0.223 |
| Social Motivation | 93 | 264 | 61.860 | 63.538 | -1.174 | 0.242 | 0.283 | 0.140 |
| Restricted and Repetitive Behavior | 93 | 264 | 66.301 | 70.640 | -3.031 | 0.003 | 0.003 | 0.366 |
| SWAN Total | 93 | 264 | 0.321 | 1.111 | -7.547 | < 0.001 | < 0.001 | 0.942 |
| Full-Scale IQ | 63 | 219 | 94.571 | 97.018 | -0.864 | 0.390 | 0.361 | 0.125 |
| CBCL Total T | 93 | 264 | 60.118 | 63.258 | -2.797 | 0.006 | 0.006 | 0.339 |
| Age (years) | 99 | 267 | 11.130 | 10.577 | 1.144 | 0.255 | 0.476 | -0.141 |
| Anxiety diagnosis (proportion) | 99 | 267 | 0.333 | 0.405 | -1.263 | 0.208 | 0.215 | 0.147 |
| Oppositional/conduct diagnosis (proportion) | 99 | 267 | 0.081 | 0.180 | -2.732 | 0.007 | 0.020 | 0.296 |
| Number of diagnoses | 99 | 267 | 2.748 | 3.790 | -5.257 | < 0.001 | < 0.001 | 0.626 |

**S6.2. Autistic trait severity**

The comparison runs opposite to the confounding account. The comorbid group scored *higher* on SRS-2 Total T than the group with autism alone (M = 70.13 against 66.85, Welch’s t = −2.58, p = 0.011, Cohen’s d = 0.31; Mann-Whitney p = 0.014).

Because SRS scores are known to be inflated by behavioral and attentional symptomatology (Hus et al., 2013), a total-score elevation in a group defined partly by attentional symptoms is not decisive on its own. We therefore examined the subscales. Two differed significantly: Social Awareness (62.01 against 66.73, p = 0.0007, d = 0.42) and Restricted and Repetitive Behavior (66.30 against 70.64, p = 0.003, d = 0.37). Social Cognition (p = 0.113), Social Communication (p = 0.060) and Social Motivation (p = 0.242) did not.

The Restricted and Repetitive Behavior subscale carries the argument. It is the SRS component least dependent on attentional and externalizing symptomatology, and its elevation in the comorbid group cannot be attributed to instrument contamination in the way a Social Awareness elevation can — noticing social cues plausibly depends on sustained attention, so that subscale is among those a co-occurring attentional diagnosis would be expected to inflate. An elevation confined to attention-loaded subscales would have been consistent with contamination rather than with greater core severity; an elevation that includes restricted and repetitive behavior is not.

SWAN Total differed as expected (0.32 against 1.11, d = 0.94). This is definitional rather than informative, since it is the variable on which the groups are constituted.

**S6.3. Other clinical dimensions**

The groups did not differ in full-scale IQ (94.57 against 97.02, p = 0.390), age (11.13 against 10.58, p = 0.255) or clinician-recorded anxiety diagnosis (33.3% against 40.5%, p = 0.208). The IQ comparison rests on a reduced sample, since Wechsler scores were available for 63 of 99 and 219 of 267 participants respectively, and is correspondingly less well powered.

The comorbid group had higher CBCL Total T-scores (60.12 against 63.26, p = 0.006, d = 0.34), a higher rate of oppositional or conduct diagnosis (8.1% against 18.0%, p = 0.007) and carried 1.04 more diagnoses on average (2.75 against 3.79, p < 0.001). The diagnostic-count difference is accounted for exactly by the ADHD diagnosis itself.

**S6.4. Direction of the potential confound**

Every clinical dimension on which the two groups differ — autistic trait severity including restricted and repetitive behavior, general psychopathology, oppositional and conduct diagnoses, and diagnostic load — is elevated in the comorbid group, and the comorbid group is the one showing the *smaller* reduction in gaze synchrony. Confounding by clinical severity would therefore be expected to work against the observed interaction rather than to produce it.

This expectation is borne out directly. Adding oppositional or conduct diagnosis and CBCL Total T-score as covariates strengthened the interaction in every context (Section S3.3, Table S9) rather than attenuating it.

**Supplementary Section S7: Stimulant Medication**

**S7.1. Availability of medication data**

Stimulant medication affects pupil dynamics and attentional engagement, and stimulant use is far more common in children with ADHD than in children without. If the comorbid group were substantially medicated on the day of assessment, pharmacological normalization could contribute to the attenuation reported in the main text.

Acute medication status was recorded with the HBN DailyMeds instrument, which logs medications active on the day of the laboratory visit. Status was available for 514 of 2476 participants (20.8%) and unknown for 1962 (79.2%) (Supplementary Table S18). Unknown status is treated throughout as unknown and never as unmedicated.

**Supplementary Table S18.** Acute stimulant status across the analytic sample, as recorded by the HBN DailyMeds instrument on the day of assessment. Percentages are of the full analytic sample; status is unknown for the majority and is not treated as unmedicated.

| Medication Status | n | % analysis sample |
| --- | --- | --- |
| Unknown | 1962 | 79.24 |
| No stimulant | 480 | 19.39 |
| Stimulant | 34 | 1.37 |

Availability varied across diagnostic groups, and not in the direction institutional prioritization of clinically diagnosed participants would predict: status was known for 36.8% of typically developing participants, 23.2% of the autism-without-ADHD group, 21.4% of the “Other” group, 20.6% of the comorbid group and 16.9% of the ADHD-without-autism group. The best-characterized group is therefore the one with no clinical diagnosis. We have not been able to establish what drove this pattern, and note it as a limitation on what the medication analyses can support.

**S7.2. Prevalence of active stimulant use**

Among participants whose status is known, 34 of 514 (6.6%) were taking a stimulant on the day of assessment; across the whole analytic sample this is 1.4% (Table S18). By group, among those tracked (Supplementary Table S19): 14.5% of the comorbid group (8 of 55), 11.3% of the ADHD-without-autism group (22 of 195), 2.6% of the “Other” group (4 of 153), and none of the autism-without-ADHD group (0 of 23) or the typically developing group (0 of 88).

**Supplementary Table S19.** Acute stimulant status by diagnostic group. The final column is computed over participants with known status only, which is a minority of each group and varies across groups from 16.9% to 36.8%.

| Diagnosis | No stimulant | Stimulant | Unknown | % stimulant of tracked |
| --- | --- | --- | --- | --- |
| TD | 88 | 0 | 151 | 0 |
| ADHD only | 173 | 22 | 961 | 11.3 |
| ASD only | 23 | 0 | 76 | 0 |
| ASD and ADHD | 47 | 8 | 212 | 14.5 |
| Other | 149 | 4 | 562 | 2.6 |

The zero in the autism-without-ADHD group rests on 23 participants of 99 and should not be read as establishing that the group was unmedicated. What the figures do establish is that among participants whose status is known, confirmed stimulant use is uncommon in absolute terms in every group, including the comorbid group that the pharmacological account would require to be substantially medicated.

**S7.3. Models adjusted for medication status**

We refitted the primary models with medication status as a three-level covariate — confirmed stimulant, confirmed no stimulant, unknown — which preserves the full sample while absorbing whatever systematic difference exists between tracked and untracked participants (Supplementary Table S20).

**Supplementary Table S20.** The 2 × 2 analysis of covariance with medication status entered as a three-level covariate (stimulant, no stimulant, unknown). Estimates are indistinguishable from the primary specification of main text Table 2.

| Metric | Term | Video | n | β | η_p_^2^ | p | q |
| --- | --- | --- | --- | --- | --- | --- | --- |
| Gaze  ISC  Magnitude | ASD | TP | 1257 | -0.785 | 0.021 | < 0.001 | < 0.001 |
|  | ASD | DM | 1390 | -0.782 | 0.022 | < 0.001 | < 0.001 |
|  | ASD | WK | 1455 | -0.598 | 0.014 | < 0.001 | < 0.001 |
|  | ASD | FF | 854 | -0.439 | 0.008 | 0.010 | 0.010 |
|  | ADHD | TP | 1257 | -0.124 | 0.002 | 0.163 | 0.327 |
|  | ADHD | DM | 1390 | -0.151 | 0.003 | 0.064 | 0.258 |
|  | ADHD | WK | 1455 | -0.065 | 0.001 | 0.407 | 0.407 |
|  | ADHD | FF | 854 | -0.108 | 0.002 | 0.259 | 0.345 |
|  | ASD × ADHD | TP | 1257 | 0.497 | 0.007 | 0.004 | 0.005 |
|  | ASD × ADHD | DM | 1390 | 0.649 | 0.012 | < 0.001 | < 0.001 |
|  | ASD × ADHD | WK | 1455 | 0.459 | 0.006 | 0.002 | 0.005 |
|  | ASD × ADHD | FF | 854 | 0.266 | 0.002 | 0.175 | 0.175 |
| Absolute  Gaze  Phase | ASD | TP | 1257 | 0.000 | 0.000 | 0.999 | 0.999 |
|  | ASD | DM | 1390 | 0.406 | 0.006 | 0.004 | 0.016 |
|  | ASD | WK | 1455 | 0.350 | 0.005 | 0.010 | 0.020 |
|  | ASD | FF | 854 | 0.064 | 0.000 | 0.712 | 0.949 |
|  | ADHD | TP | 1257 | 0.076 | 0.001 | 0.401 | 0.624 |
|  | ADHD | DM | 1390 | 0.059 | 0.000 | 0.468 | 0.624 |
|  | ADHD | WK | 1455 | 0.134 | 0.002 | 0.098 | 0.390 |
|  | ADHD | FF | 854 | -0.030 | 0.000 | 0.756 | 0.756 |
|  | ASD × ADHD | TP | 1257 | 0.037 | 0.000 | 0.832 | 0.832 |
|  | ASD × ADHD | DM | 1390 | -0.388 | 0.004 | 0.015 | 0.060 |
|  | ASD × ADHD | WK | 1455 | -0.243 | 0.002 | 0.115 | 0.230 |
|  | ASD × ADHD | FF | 854 | -0.089 | 0.000 | 0.657 | 0.832 |
| Pupil  ISC | ASD | TP | 757 | -0.811 | 0.024 | < 0.001 | < 0.001 |
|  | ASD | DM | 827 | -0.456 | 0.008 | 0.010 | 0.021 |
|  | ASD | WK | 917 | -0.347 | 0.005 | 0.041 | 0.055 |
|  | ASD | FF | 401 | -0.146 | 0.001 | 0.575 | 0.575 |
|  | ADHD | TP | 757 | -0.065 | 0.000 | 0.591 | 0.862 |
|  | ADHD | DM | 827 | 0.019 | 0.000 | 0.862 | 0.862 |
|  | ADHD | WK | 917 | -0.073 | 0.001 | 0.486 | 0.862 |
|  | ADHD | FF | 401 | 0.060 | 0.001 | 0.674 | 0.862 |
|  | ASD × ADHD | TP | 757 | 0.563 | 0.009 | 0.010 | 0.040 |
|  | ASD × ADHD | DM | 827 | 0.083 | 0.000 | 0.681 | 0.822 |
|  | ASD × ADHD | WK | 917 | 0.284 | 0.002 | 0.143 | 0.286 |
|  | ASD × ADHD | FF | 401 | 0.068 | 0.000 | 0.822 | 0.822 |

Estimates were indistinguishable from the primary specification. The ASD × ADHD interaction on gaze synchrony was +0.497 (q = 0.005) in *The Present*, +0.649 (q = 0.0002) in *Despicable Me* and +0.459 (q = 0.005) in *Diary of a Wimpy Kid*, against +0.497, +0.649 and +0.459 in the primary models — agreement to three decimal places. In *Fun with Fractals* the estimate moved by 0.011 SD (+0.266 against +0.277) and remained non-significant. Autism main effects likewise moved by at most 0.012 SD in any context, and the pupil and phase results were similarly unchanged.

**S7.4. What this establishes**

A missing-indicator covariate does not recover an unbiased estimate of a medication effect when status is not missing at random, and status is plainly not missing at random here (Section S7.1). That is not the question it is being asked to answer. The question is whether the diagnostic interaction is sensitive to medication variance among participants for whom medication variance can be measured, and it is not: the estimates are essentially unchanged.

This establishes that stimulant exposure does not account for the interaction among participants whose status is documented. It does not exclude pharmacological effects among the 79.2% whose status is unknown. We therefore report medication as a sensitivity analysis that the result survives, rather than as having ruled out a pharmacological explanation.
